# ELISL: Early-Late Integrated Synthetic Lethality Prediction in Cancer

**DOI:** 10.1101/2022.09.19.508413

**Authors:** Yasin Tepeli, Colm Seale, Joana Gonçalves

## Abstract

Anti-cancer therapies based on synthetic lethality (SL) exploit tumor vulnerabilities for treatment with reduced side effects. Since simultaneous loss-of-function of SL genes causes cell death, tumors with known gene disruptions can be treated by targeting SL partners. Computational selection of promising SL candidates amongst all gene combinations is key to expedite experimental screening. However, current SL prediction models: (i) only use tissue type-specific molecular data, which can be scarce/noisy, limiting performance for some cancers; and (ii) often rely on shared SL patterns across genes, showing sensitivity to prevalent gene selection bias. We propose ELISL, Early-Late Integrated models for SL prediction using forest ensembles. ELISL models ignore shared SL patterns, and integrate context-specific data from cancer cell lines or tumor tissue with context-free functional associations derived from protein sequence. ELISL outperformed existing methods and was more robust to selection bias in 8 cancer types, with prominent contribution from sequence. We found better survival for patients whose tumors carried simultaneous mutations in a BRCA gene together with an ELISL-predicted SL gene from the HH, FGF, or WNT families. ELISL thus arises as a promising strategy to discover SL interactions with therapeutic potential.

## Introduction

Targeted anti-cancer therapy can be preferred to conventional chemo and radiotherapy, since it reduces side effects by capitalizing on tumour-specific molecular changes to selectively kill tumour cells. However, direct drug targeting may be prevented by alterations affecting the drug target, such as loss of function mutations, amplification, or overexpression [1, 2]. A promising alternative explores synthetic lethality (SL) between a group of genes, whereby simultaneous dysfunction of all genes in the SL group causes cell death while disruption of only a subset of those SL genes is non-lethal [3]. Tumours with a known dysfunctional gene can then be treated by targeting SL partner genes.

The clinical approval of PARP inhibitor drugs for treatment of BRCA deficient tumours has confirmed the promise of SL-based therapies [4, 5], yet the number of validated SL relations remains scarce. The main reason for this is that identifying SL interactions requires costly and laborious molecular perturbation experiments [678910], making it infeasible to perform exhaustive screening of all gene combinations in many samples or conditions of interest. Computational SL prediction is therefore crucially needed to prioritize SL candidates for experimental follow-up.

Existing SL prediction methods fall into two main categories, statistical approches and machine learning (ML) models. Statistical methods such as DAISY [11], BiSep [12], and ISLE [13] select SL pairs by imposing thresholds on statistical properties associated with SL, such as mutual exclusivity of mutations in both genes gene coexpression, or changes in cell dependency on a gene. Statistical models are intuitive. However, they are unable to capture more complex relationships underlying SL interactions and tend to underperform compared to ML-based models [14].

The ML-based models can be further split into SL-topology and feature-based approaches. SL-topology methods represent existing SL interactions as networks, with nodes denoting genes and edges representing pairwise SL interactions. SL-topology methods use this SL network to identify shared SL patterns across genes and infer new SL interactions using matrix factorization (pca-gCMF [15], GRSMF [16], and SL2MF [17]) or graph-based methods (DDGCN [18] and GCATSL [19]). The dependence of SL-topology methods on existing SL interactions means that (i) their prediction scope is often limited to genes with known SL partners, and (ii) they are more suitable for transferring known SL interactions between genes with similar SL profiles than for *de novo* SL discovery. The performance of SL-topology methods is also heavily influenced by connectivity, but typically SL data covers only a small fraction of all possible genes and interactions, resulting in rather sparse SL networks. Additionally, existing SL data has a prevalent selection bias towards functionally related genes, showing similar SL profiles with other genes that SL-topology methods are designed to exploit. However, SL interactions among a few related genes do not generalize to most unseen genes, making SL-topology methods sensitive to gene selection bias [14].

Feature-based ML models are built with supervised learning algorithms using features derived from molecular profiles (DiscoverSL [20], EXP2SL [21], Lu [22], and SBSL [14]). In this way, SL-feature ML models are able to incorporate complex rules underlying SL interactions while remaining less sensitive to selection bias. Most feature-based methods rely on (regularized) logistic regression or random forests to predict SL based on multiple omics features [20,22,14]. Alternatively, EXP2SL uses a neural network to learn from a fixed set of genes and their expression in cancer cell lines [21]. Common to all existing SL-feature models is that they focus only on context-specific molecular data for a tissue type of interest: for lung cancer, this could be omics data of lung cancer cell lines and lung tumour tissue. While context-specific data is valuable for SL prediction, it may be unavailable or difficult to obtain for some (rarer) cancer types, limiting the ability to learn useful SL prediction models.

We argue that broader context-free measures of functional similarity between genes may have added value for SL prediction. The idea is that genes of similar function have more intertwined or redundant activity, making it more likely that a (cancer) cell would depend on their joint loss of function for its survival [23]. Homology of amino acid sequences and similarity of protein-protein interactions (PPI) have been used successfully as proxies for functional similarity in areas such as protein function prediction [24,25,26], and are therefore good candidate metrics. Sequence homology, in particular, is very appealing since amino acid sequences are readily available and (nearly) complete.

We propose *Early-Late Integrated Synthetic Lethality* (ELISL) prediction models, the first to integrate context-free and context-specific omics features to predict SL for pairs of genes. Context-free features in ELISL express the difference between the vectorized embeddings of the genes in the pair, based on their amino acid sequences or PPIs. Context-specific features are stratified per tissue and sample type. For cancer cell lines, ELISL considers changes in dependency to perturbation on one gene when the other gene is altered based on mutation, expression, or copy-number data. For tumour and healthy tissues, expression changes of one gene based on alterations in the other, as well as coexpression, are included. For patient tumours, ELISL also considers changes in patient survival between groups with or without simultaneous mutations in both genes. To effectively learn from both low and high dimensional data sources, across sparser and denser representations, ELISL models rely on a combination of early and late integration strategies using a collection of decision tree ensembles.

## Results and Discussion

The Early Late Integrated Synthetic Lethality (ELISL) prediction framework leverages context-free and context-specific functional relationships to predict synthetic lethality of pairs of genes. To achieve this goal, ELISL uses a combination of early and late integration to jointly learn from context-free similarity of protein sequences and PPIs, and from context-specific statistics derived from molecular and clinical data of cancer cell lines and human tissue.

ELISL aggregates six regularized forest ensemble models (Fig. 1a). One model learns from all features, enabling early integration and feature interactions across data sources. Each of the remaining five models learns from a different context-free or context-specific source to enable late integration and balance between low and high dimensional data. Two cell line models explore the relationship between dependency of cells upon perturbation of one gene (DepMap [27, 28]) and presence of alterations in the other gene (CCLE [29,30,31,32]), based on mutations and expression (4 features). Alterations are generally defined by the presence of one of the following: non-silent mutations, expression z-scores larger than 1.96 or smaller than -1.96, or discrete copy number values of 2 (amplified) or -2 (deleted). Additionally, a tissue model focuses on tissue omics and clinical data from healthy donors (GTEx, [33]) and tumour patients (TCGA, [31,32,34]). This model considers: average expression of one gene in samples with or without alterations in the other gene (4 features); coexpression of the gene pair in tumour, normal, or healthy donor samples (Pearson’s correlation and p-value, 6 features); correlation of copy number values in tumour samples (Spearman’s correlation and p-value, 2 features); and change in patient survival between groups with and without simultaneous mutations in both genes (Cox regression test and p-value, 4 features). Finally, two context-free models learn from embeddings of amino acid sequences from UniProt [35] using SeqVec [36] (1024 features), or nodes in a network of curated PPIs from STRING [37] using Node2Vec [38] (64 features). The feature vector of a gene pair is defined as the element-wise absolute difference between the embedding vectors of the two genes.

**Figure 1.**
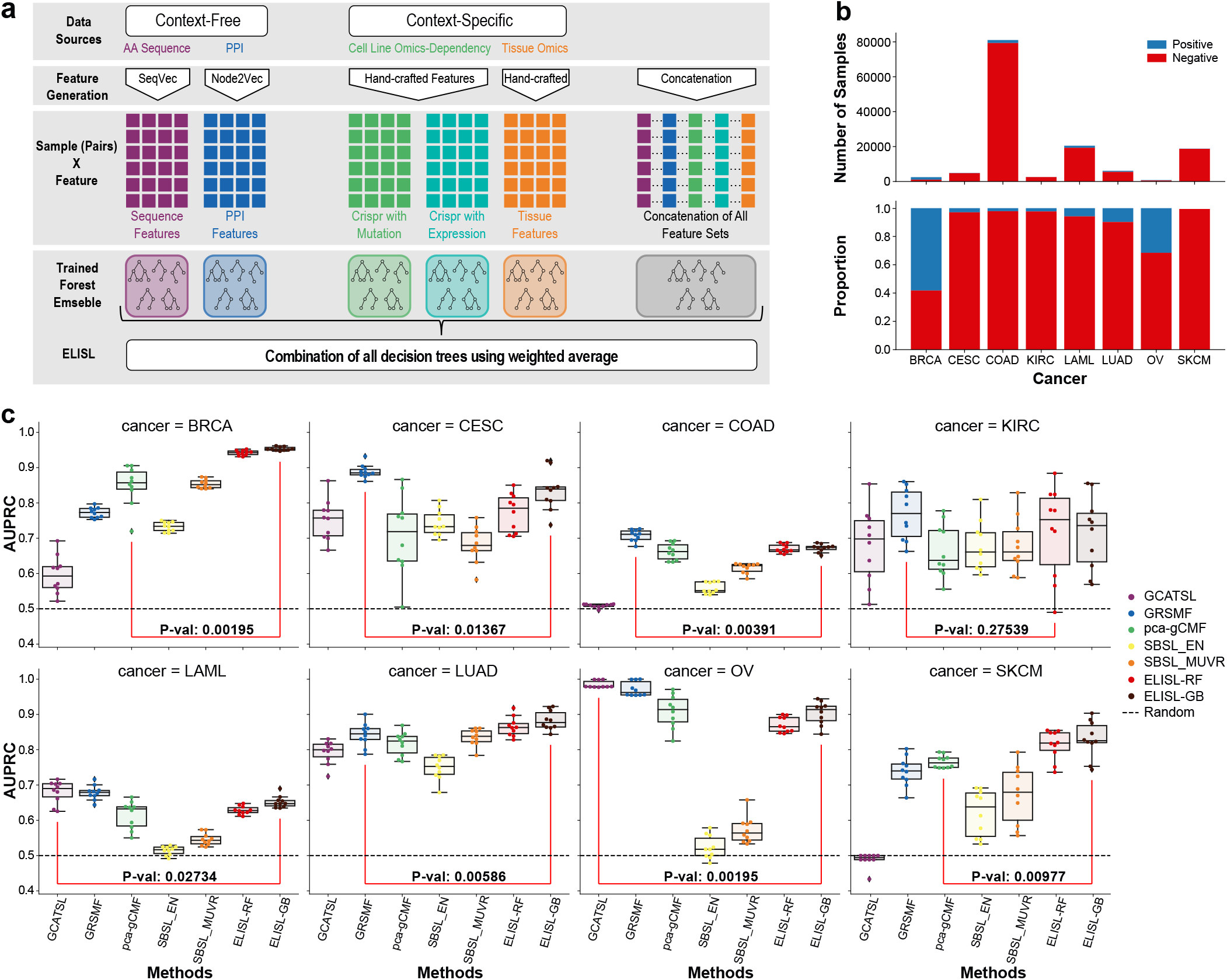
ELISL framework, SL label imbalance, and within cancer prediction performance. **a**, The ELISL framework: six forest ensemble models learn separately from two context-free sources, three context-specific sources, and all data sources together. Context-free sources, amino acid sequence and PPI. Context-specific sources include: cell line dependency and mutation; cell line dependency and expression; tissue mutation, expression, copy number, and patient survival. Features for context-free sources express sequence and PPI similarity based on embeddings, while features for context-specific sources denote various statistical measures based on molecular profiles. The six models are ensembled using weighted average to obtain the final prediction probability for each gene pair. **b**, Number and ratio of positive and negative samples in the train set for each cancer type. **c**, Prediction performance (AUPRC) of SL prediction methods per cancer type over 10 runs using independently drawn 80/20 train/test splits (same cancer). Method categories: matrix factorization and graph-based (GCATSL, GRSMF, pca-gCMF); supervised learning, including existing models (SBSL-EN/MUVR), and our proposed ELISL models (ELISL-RF/GB). Red lines compare the best ELISL model with the best among the other models. P-val shows the significance of the difference in performance between the models over 10 runs, using a two-sided Wilcoxon signed rank test.

We built two types of ELISL models using random forests (ELISL-RF) or gradient-boosted decision trees (ELISL-GB). Models were trained on experimentally validated SL labels made available by previous studies for different cancer types (DiscoverSL [20], ISLE [13], Exp2SL [21], Lu et al. [22]). Since SL labels were highly imbalanced, we used random undersampling of the majority class (non-SL or negative) to mitigate learning bias. (Fig. 1b). An ELISL prediction denotes a probability that a pair of genes is SL for a given tissue type, calculated as a weighted average of the predictions of its six underlying models (see Methods). Prediction performance on test sets was summarized using area under the precision-recall curve (AUPRC) or ROC curve (AUROC), and Matthews coefficient (MCC).

### Cancer-specific synthetic lethality predictions

To evaluate the ELISL models, we first assessed their ability to generalize within each cancer type, for eight distinct cancer types. We compared ELISL-RF and ELISL-GB to five other recently published ML models with high performances in their categories, namely: pca-gCMF [15], GRSMF [16], and GCATSL [19] as matrix factorization and graph-based methods, and SBSL-MUVR and SBSL-EN [14] as supervised ML models. Models were evaluated per cancer type using ten independent 80/20 splits of the SL labelled gene pairs (Methods, Supplementary Section 2.3).

Supervised ELISL models outperformed all other methods in breast (BRCA), lung (LUAD), and skin (SKCM) cancers, while graph-based GRSMF or GCATSL outperformed the other methods in cervix (CESC), colon (COAD), leukemia (LAML), and ovarian (OV) cancers (AUPRC Fig. 1c, AUROC and MCC Supplementary Fig. S3). For kidney (KIRC) cancer, all methods showed high variance, and there was no clear best performing model. ELISL models achieved significantly higher performance than the second best model for BRCA, LUAD, and SKCM (Wilcoxon *p* ≤ 0.01). In addition, ELISL remained competitive with graph-based methods as the second best (CESC, COAD), or third best model (LAML, OV). In contrast, the two graph-based models were in the bottom three for BRCA, and GCATSL failed to learn useful models for both COAD and SKCM.

Matrix factorization and graph-based models had strikingly high performances in OV. This is consistent with our previous finding that SL-topology methods thrive on OV due to selection bias [14]. We also reported that SL labelled pairs for OV span a limited set of functionally related genes, where the bias was visually apparent through the presence of consistent SL patterns across the genes.

### Robustness of synthetic lethality predictions to gene selection bias

We recently reported that the performance of SL prediction models can be overestimated due to gene selection bias in SL labels, particularly when predictions are driven by shared patterns of existing SL interactions [14]. To assess the impact of gene selection bias on ELISL and the remaining five methods, we performed two experiments where the sets of gene pairs used for training (train set) and evaluating (test set) followed different gene selection biases.

#### Double gene holdout (induced differences in gene bias)

For the double gene holdout experiment, we ensured that the train and test sets followed different gene selection biases. To do this, we constructed ten matched train and test sets such that there were no genes shared between each train set and its matched test set. This differs from the original experiment where matched train/test sets were allowed to follow similar gene selection bias, by preventing overlap only in terms of gene pairs and not individual genes. All methods were evaluated in four cancer types, BRCA, CESC, LUAD, and OV. We excluded KIRC and SKCM due to the limited number of gene pairs, and COAD and LAML due to generally poor performances in the original experiment (Fig. 1).

Using double gene holdout, the performances of all models decreased significantly for all cancer types (AUPRC Fig. 2a, AUROC and MCC Supplementary Fig. S4), which could partly be caused by the considerable reduction imposed by the train/test set construction on the number of gene pairs available for training (Supplementary Table S5). For BRCA, the two ELISL models performed the best (median AUPRC ELISL-RF 0.67, ELISL-GB 0.69), while matrix factorization and graph-based methods dropped to nearly random (Fig. 2a, top left). For CESC, GRSMF had performed significantly better than ELISL in the original single cancer experiment, but the difference between the methods dissipated using double gene holdout (Fig. 2a, top right). For LUAD, most methods struggled with double gene holdout Fig. 2a, bottom left). However, the supervised ML models SBSL and ELISL retained above random performances, with ELISL-RF achieving the best median AUPRC (0.59). For OV, we saw the largest decrease in performance using double holdout compared to the original experiment, which was expected given the known bias in SL labels. ELISL-RF and GRSMF performed the best in OV using double gene holdout, while the SBSL models maintained their originally modest performances (Fig. 2a, bottom right). The GCATSL method, best of the original experiment (0.98 median AUPRC), dropped to near random performance with double holdout (0.5 median AUPRC).

**Figure 2.**
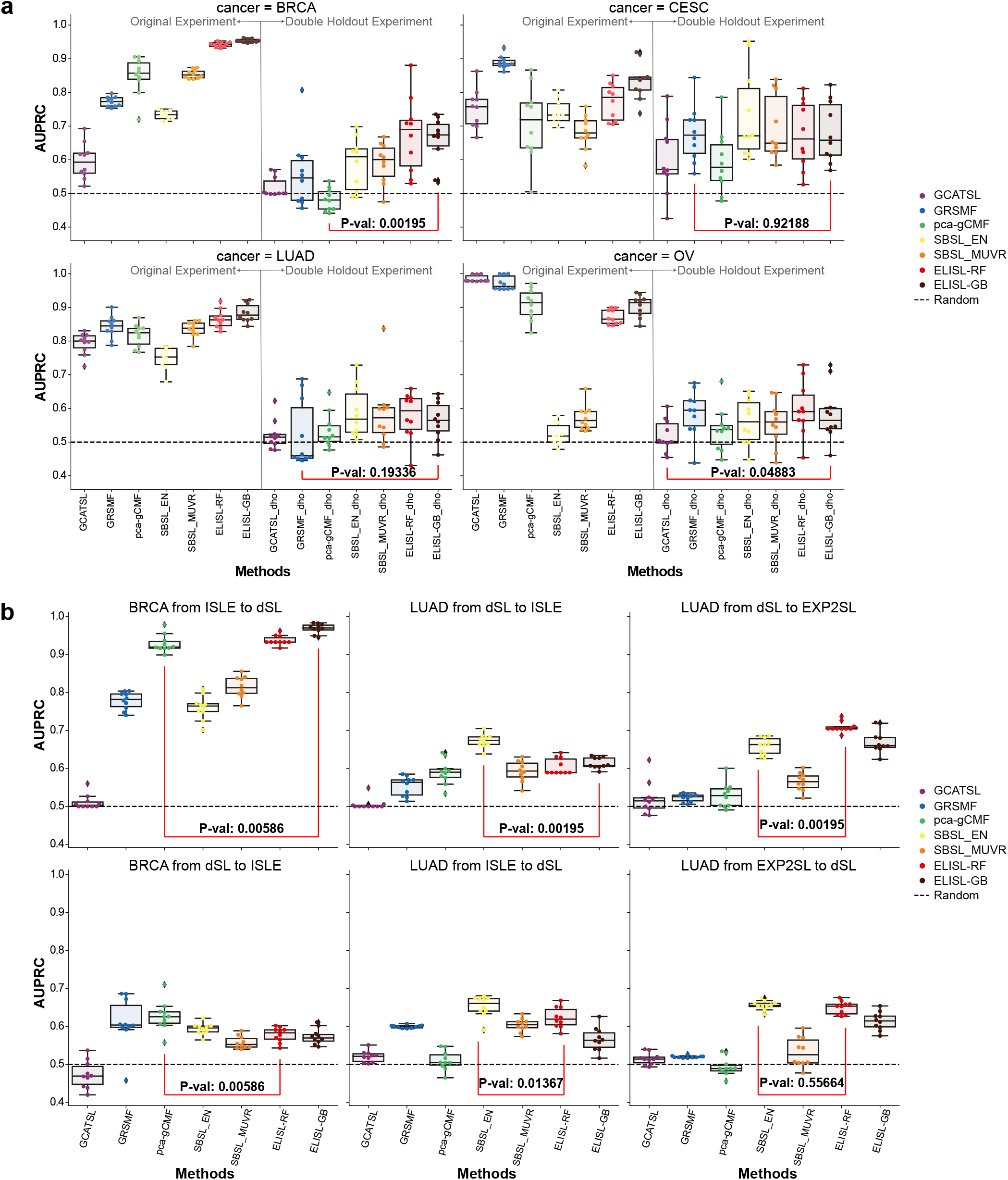
Impact of gene selection bias on the performance of SL prediction models. **a**, Double gene holdout experiment to assess the impact of induced differences in gene selection bias between train and test set. Performance (AUPRC) per cancer type and for 10 runs with independently drawn 80/20 train/test splits, where each pair of train and test sets does not share any genes. For each cancer, the left side of the plot reports the original performance (same as in Fig. 1c), and the right side shows the performance using double gene holdout. P-val denotes the significance of the difference between the double holdout performances of the two models that performed best in the original experiment. **b**, Cross-SL dataset experiment to assess the impact of inherent differences in gene selection bias between different datasets of SL labels (ISLE, DiscoverSL, and EXP2SL). Performance (AUPRC) is reported for models trained using labels from one SL dataset and evaluating on another SL dataset, considering the combinations of cancer type (BRCA, LUAD) and SL datasets with sufficient number of samples.

Overall, the supervised ML models SBSL and ELISL did better using double gene holdout. Matrix factorization and graph-based methods delivered inconsistent performances across cancer types, indicating that they are more sensitive to gene selection bias. ELISL models outperformed the other methods in BRCA and LUAD, and were comparable to the best performing models in CESC and OV. Among the two ELISL models, ELISL-RF seemed slightly more successful in the double gene holdout experiment compared to ELISL-GB.

#### Cross-SL label prediction (inherent differences in gene bias)

Since the double holdout scenario is an extreme case designed by us, we set out to evaluate the SL prediction models in a setting with naturally occurring differences in gene selection between train and test sets. To do this, we trained the models using SL labelled pairs from one source and tested them on labelled pairs from another source for the same cancer type. We used the following combinations of SL labelled pairs, which yielded a sufficient number of samples for evaluation (between 78 and 1146, Supplementary Table S6): for BRCA, training on ISLE labelled pairs and testing on DiscoverSL; for LUAD, training on DiscoverSL labels and testing on Exp2SL or Lu et al. labels. We also tested the reverse combinations, but results were less reliable due to the smaller number of training pairs they had in comparison (Supplementary Table S6).

ELISL models outperformed the other methods when training on ISLE and predicting on DiscoverSL for BRCA and training on DiscoverSL and predicting on Exp2SL for LUAD (AUPRC Fig. 2b, (AUROC Supplementary Fig. S5)). For the remaining experiments on LUAD, one of the ELISL models was always amongst the top 2 performing methods, where the linear SBSL-EN model typically took the lead. The only experiment in which ELISL showed poor performance compared to other methods was when training on dSL and predicting on ISLE for BRCA. Overall, supervised ML models emerged again as the most robust to selection bias, just like in the gene holdout experiments. The SBSL-EN and ELISL-RF models performed reasonably in most experiments, while GRSMF was the best among the matrix factorization and graph-based methods.

#### ELISL-RF prediction of synthetic lethality across cancer types

Synthetic lethal interactions may occur in multiple cancer types. For instance, PARP inhibitor drugs have been approved for treatment of breast, ovarian, prostate[39], and pancreatic[40] cancers carrying BRCA deficiencies, suggesting that the SL relationship BRCA-PARP is not cancer-specific[41]. This suggests that there could be some benefit in leveraging successful models trained on cancer types with sufficient data (BRCA, LUAD, OV) to predict SL in other cancer types for which samples are either not available or difficult to obtain (CESC, KIRC, and SKCM). We investigated this by evaluating ELISL-RF models trained per cancer type against each of the remaining cancer types.

The success of cross-cancer SL predictions was modest for most pairwise combinations of cancer types, to which the quality and biases of the labels could have contributed as well (AUPRC Fig. 3a, (AUROC Supplementary Fig. S7)). Nevertheless, we also saw some promising results. For the prediction of CESC pairs, the LUAD-trained model performed better than the CESC-trained model itself (0.85 vs. 0.77 mean AUPRC). Models trained on COAD or KIRC also achieved reasonable performances in CESC (0.69 and 0.71 mean AUPRC respectively). For SL prediction in KIRC, the best model was trained using KIRC labelled pairs (0.72 mean AUPRC). However, models trained on CESC, BRCA, LUAD showed reasonable performances as well (>0.63 mean AUPRC). Additionally, promising results of training on KIRC and predicting on BRCA, training on LAML and predicting on CESC, and training on LAML and predicting on LUAD (all > 0.65 mean AUPRC) might indicate common SL pairs across the cancer types.

**Figure 3.**
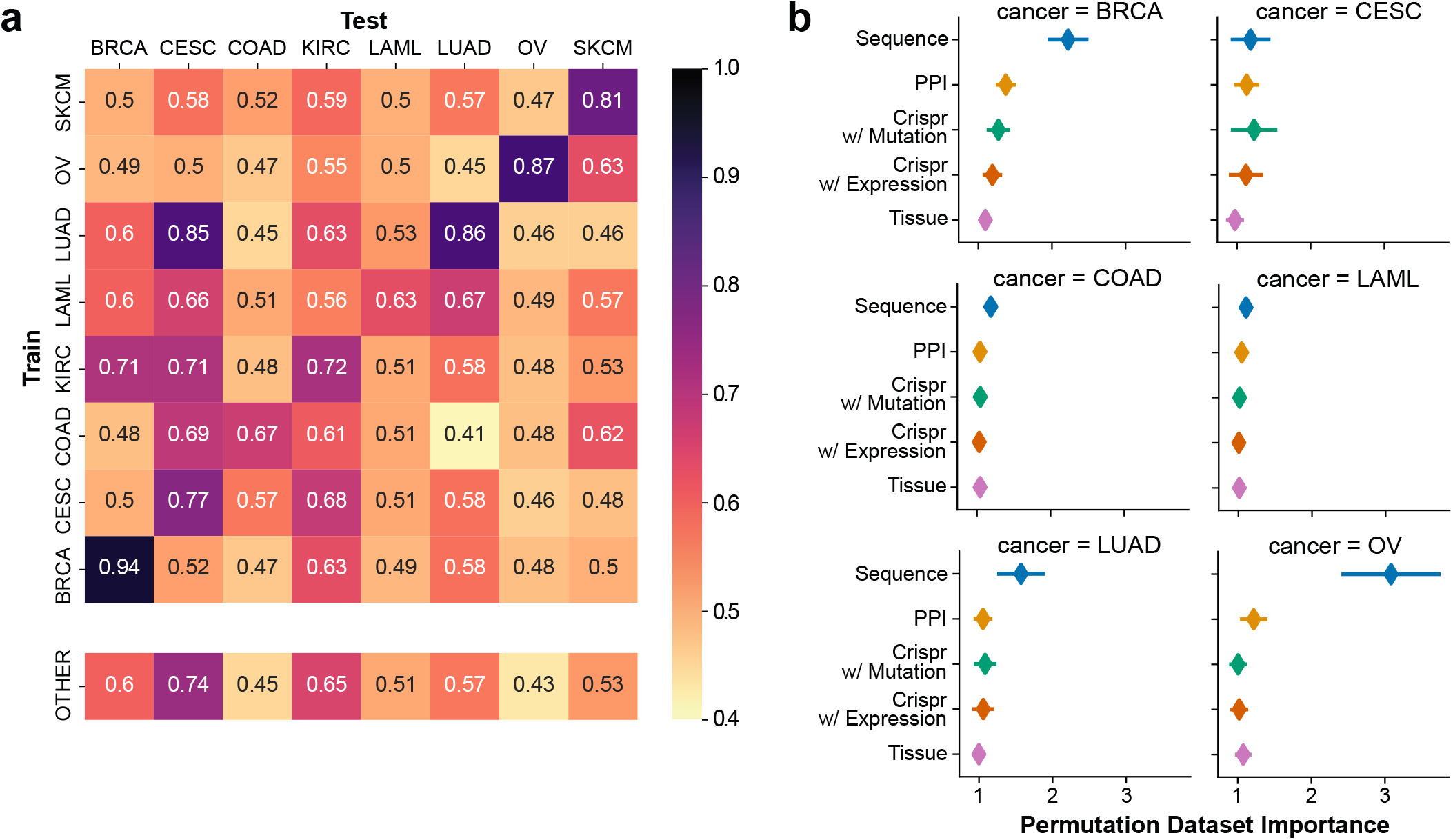
ELISL-RF SL prediction within/across cancer types and feature contribution. **a**, Performance of cancer-specific models and pan-cancer models, measured as average AUPRC over 10 runs using independently drawn 80/20 train/test splits. For cancer-specific models, presented in a matrix, the diagonal reports prediction performance within the same cancer type, and the remaining cells show performance for predicting on other cancer types. Pan-cancer model performances are reported in a separate row at the bottom, where models are trained on all other cancer types except the one the model is supposed to predict on. Rows denote the cancer type used for training, columns indicate the cancer type used for prediction and evaluation. **b**, Contribution of each data source to the predictions of the ELISL-RF model within the same cancer type, whose performances can be seen in the matrix diagonal of Fig. 3a. Calculated using permutation feature importance, with importance defined as the ratio of errors (1-AUPRC) between predictions using the permuted vs original test samples.

Inspired by the results, we investigated if models learnt using SL labels from multiple cancer types (pan-cancer) would provide any benefit compared to cross-cancer predictions. For every cancer type *T*, we trained models using labels from all other cancer types (except *T*), and then evaluated the predictions for labelled pairs in *T* (see Methods). Pan-cancer models showed promising performance for CESC (0.74 mean AUPRC) and reasonable results for KIRC (0.65; Fig. 4a, top row). Performances of pan-cancer models were not better than those of cancer-specific and crosscancer models, indicating that prior selection of relevant cancer types could be needed to achieve better performances and effectively enable pan-cancer models to predict SL for cancer types with limited sample sizes.

**Figure 4.**
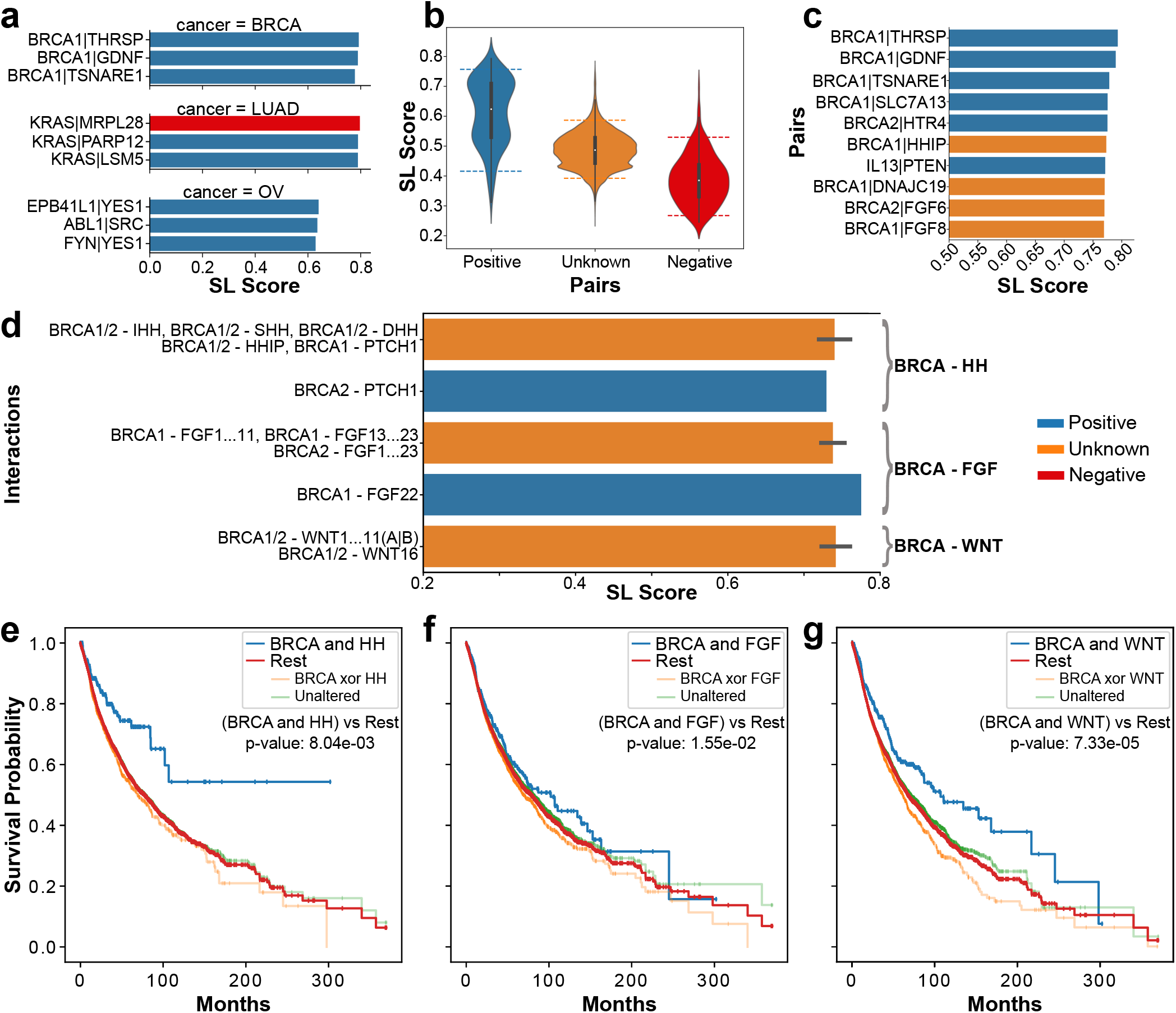
Analysis of top SL gene pairs predicted by ELISL-RF. **a**, Top 3 pairs ranked by SL prediction score for the test sets of BRCA, LUAD, OV. Figures **b** to **g** show results for prediction of unknown gene pairs (not in the test) using the ELISL-RF trained on BRCA data. **b**, Distribution of SL scores for positive, unknown, and negative pairs. Positive and negative pairs come from the BRCA test set, unknown pairs are generated using genes found in cancer and DNA repair pathways from KEGG, Reactome, and PID. The dashed lines denote the 5% and 95% percentile. **c**, SL prediction scores of the top 10 pairs across the BRCA test set and generated unknown set using ELISL-RF. **d**, SL prediction scores of pairs between hedgehog (HH) family members and BRCA1/2; between fibroblast growth factor (FGF) family members and BRCA1/2; and between WNT family members and BRCA1/2. The length of each bar denotes the average SL score and the length of the black line at the top of each bar represents the standard deviation for the set of pairs of interest. Figures **e, f**, and **g** show differences in survival between groups of patient tumour samples with and without simultaneous mutations in both genes of a gene pair. We consider pairs between BRCA genes and members of one of three families: **(e)** HH, **(f)** FGF, and **(g)** WNT. The plots show Kaplan-Meier curves and p-values for a Cox proportional hazards (PH) model of survival time considering the presence or absence of simultaneous mutations in both genes, and adjusted for age, sex, and cancer type.

### Contribution of context-free and -specific features to ELISL-RF models

As the ELISL-RF model integrates both context-free and context-specific features, we sought to assess the contribution of different data types to the predictions of ELISL-RF. We used a variation of permutation feature importance [42], where we permuted all features associated with each data type together. Importance was defined as the ratio of errors (1-AUPRC) of ELISL-RF predictions for each test set using the permuted vs original features (see Methods). The feature set derived from amino acid sequence data stood out as the most important in five cancer types (BRCA, COAD, LAML, LUAD, and OV), and the second most important in CESC (average importance: sequence 1.18, dependency with mutation 1.23).We note that importance values were more prominent for BRCA, CESC, LUAD, and OV because the performance of ELISL-RF was higher for these cancer types (between 0.77 and 0.94 mean AUPRC) compared to COAD and LAML (0.67 and 0.63). High performance means low errors, which can result in larger ratios (importances) for small changes in performance. Beyond sequence, PPI and the interaction of CRISPR dependency and mutation were the second most important feature sets. Ultimately, all data sources contributed to the ELISL-RF model (average importance*>* 1) in at least two cancer types. The variation in importance of each feature set across cancer types suggests that the integration of different omics data is beneficial for cross-cancer SL prediction. We also checked if the high-dimensionality of sequence embeddings influenced ELISL-RF, but using embedding sizes between 32 and 1024 led to comparable performance (Supplementary Fig. S3).

### Potential of unknown SL gene pairs predicted by ELISL-RF models

#### Predictions for labelled pairs

To further assess the potential of ELISL-RF models, we analyzed the top gene pairs ranked by prediction probability in BRCA, LUAD, and OV. All top 3 gene pairs for BRCA and OV were labelled as SL (Fig. 4a). In fact, the top 82 pairs for BRCA and top 16 pairs for OV had positive labels, confirming that ELISL-RF can recover known SL interactions. For LUAD, we counted six SL and four non-SL pairs amongst the top 10 predictions (Supplementary Table S8). Notably, the highest ranked gene pair in LUAD, KRAS-MRPL28 had a non-SL label. However, another study reported that disruption of *MRPL28* was lethal in *KRAS* -mutant cancer cell lines [43]. The finding was for colorectal cell lines, but lung cancer may share underlying mechanisms. For instance, KRAS-mutations are frequent in both lung and colorectal cancers, and colorectal cancers often metastasize to lung[44, 45]. We therefore do not discard the possibility that KRAS-MRPL28 could be mislabelled for LUAD.

#### Predictions for unknown pairs

In a separate experiment, we used ELISL-RF to make predictions for unknown gene pairs. We focused only on BRCA, since it was the cancer for which ELISL-RF models achieved the highest performance. To keep the number of unknown gene pairs manageable, we generated all pairwise combinations involving only genes present in cancer and DNA repair pathways from KEGG, Reactome, and PID (Supplementary Table S10). Overall, ELISL-RF assigned higher SL prediction scores or probabilities to pairs with known SL labels (median 0.62), and lower scores to pairs with known non-SL labels (median 0.39), as expected (Fig. 4b). The distribution of SL prediction scores for unknown pairs showed no particular tendency (median 0.49), showing that generated pairs were not biased regarding SL status.

There were three unknown gene pairs amongst the 10 pairs with the highest SL prediction scores: BRCA1-HHIP, BRCA2-FGF6, BRCA1-FGF8 (Fig. 4c). We looked into the functional roles of the genes and gene families involved in these pairs for additional insight, and investigated if the ELISL-RF predicted gene pairs have prognostic potential. Concerning the BRCA1-HHIP interaction, the hedgehog interacting protein (HHIP) binds to all three hedgehog family members (IHH, SHH, DHH) with affinity to the PTCH1 receptor, and regulates the hedgehog (HH) signaling pathway [46,47,48]. The HH pathway is synthetic lethal with the PI3K/AKT/mTOR pathway in rhabdomyosarcoma [49], and the inhibition of PI3K is known to strengthen BRCA-PARP synthetic lethality in BRCA1-deficient breast cancer [50]. We thus reason that the HHIP gene or HH family could be a synthetic lethal partner for BRCA1/2. Notably, the BRCA2-PTCH1 pair had a positive SL label [51], and all pairs between BRCA genes and HH family members yielded high prediction scores (>0.7, Fig. 4d). Additionally, analysis of patient tumour samples from TCGA showed that patients whose tumours carried mutations in both a BRCA gene (BRCA1 or BRCA2) and a HH gene (IHH, SHH, DHH, PTCH1) had significantly better survival than the rest (Cox p-value 8.04 *×* 10^*−*3^).

We assessed the BRCA2-FGF6 and BRCA1-FGF8 pairs together, since they both denote interactions between a BRCA gene (BRCA1 or BRCA2) and a member of the FGF family (FGF1 to FGF23).The fibroblast growth factor (FGF) family regulates cell differentiation and proliferation, and takes part in cancer pathogenesis [52]. Notably, the BRCA1-FGF12 pair had a positive SL interaction label (Fig. 4e), and all pairs between BRCA and FGF family members had prediction scores higher than 0.7 (Fig. 4d). The median survival time for patients whose tumours had mutations in both BRCA1/2 and FGF genes (FGF1 to FGF23) was 23 months larger than for the rest of the patients, with a Cox p-value of 1.55 × 10^*−*2^ (Fig. 4f, Supplementary Table S11).

The top 5% of gene pairs, with SL prediction scores larger than 7.57, included 8 interactions between BRCA genes and members of the WNT family (8 out of 38 or ∼ 21%, Fig. 4d and Supplementary Table S9). The WNT pathway regulates various processes including cell fate determination [53, 54]. Inhibition of WNT signaling has been found to induce a BRCA-like state, making cells vulnerable to PARP inhibition [55], which could suggest SL interactions between WNT, BRCA, and PARP. We found that patients with tumours yielding mutations in both BRCA and WNT genes had a significantly better survival probability than the rest (Cox p-value 7.35 × 10^*−*5^, Fig. 4g).

Overall, the fact that simultaneous mutations in genes from families involved in highly ranked pairs lead to differences in patient survival time indicates that ELISL-RF is able to prioritize promising SL relationships.

## Conclusion

In this work we proposed ELISL, ensemble models that leverage functional relationships between genes to predict synthetic lethality in cancer. To our knowledge, ELISL models are the first to learn from context-free (sequence) data denoting broad functional association, in addition to context-specific molecular data from cancer patient tissue or cell lines. Crucially, we consider (amino acid) sequence data for SL prediction, under the premise that sequence homology is a known proxy for related function and could thus be predictive of SL interaction. With ELISL, we introduce an early-late integrated learning strategy using forest ensembles to enable effective learning from high-dimensional sequence embeddings and more conventional features (e.g. based on gene dependency and mutation data).

ELISL models outperformed competing SL prediction methods, both with similar and different gene selection bias between train and test sets, emerging as the most accurate and robust models overall. Regardless, effective learning from gene selection bias remains a fundamental challenge that merits further research. In contrast to ELISL, some matrix factorization and graph-based models (GRSMF, pca-gCMF) performed well when train and test set followed similar gene selection bias, but struggled to make useful predictions under varying bias as we had also seen in our previous work [14]. Other feature-based models, SBSL, exhibited inconsistent performances across cancer types, exposing the weakness of relying only on context-specific features, which can be sparser and noisier for some cancers, especially rarer types with smaller numbers of available patient tissue samples.

Notably, amino acid sequence features provided by far the largest contribution to the predictions of ELISL models across cancer types, suggesting that they are indeed valuable for predicting SL interactions. Sequence was therefore also largely responsible for the advantage of ELISL models compared to our previous supervised learning approaches context-specific features (SBSL). This finding is significant, since the use of sequence embeddings allows models to be less dependent on context-specific features like gene dependencies, which are exclusively available for cellular models and may not directly translate to patient tumours.

Predicting across cancer types revealed challenging, but it was encouraging to see that ELISL models trained on colon, kidney, or lung cancer performed reasonably well on cervix cancer. Cross-cancer prediction should improve as more high quality, less biased, SL data becomes available. Nevertheless, the few successful cases point to the existence of multiple SL interactions shared between cancer types, which could benefit a larger number of patients. Using ELISL to make predictions for unknown gene pairs, we proposed three promising interactions between BRCA genes and members of the HH, FGF and WNT families. Survival analysis showed that simultaneous mutations in both groups of genes involved in these pairs coincided with longer median patient survival times, confirming that ELISL has the ability to prioritize SL interactions with effective therapeutic potential.

## Methods

### Data collection and feature generation

We generated a featurized representation for every gene pair used to learn and evaluate ELISL models. Two categories of features were derived from molecular data: context-free relationships between genes based on sequence or PPI, and context-specific features and interactions based on cell line and tissue omics.

#### Amino acid sequence

We retrieved reviewed amino acid sequences from UniProt[35], and used the SeqVec pretrained model [36] to extract a 1024-dimensional embedding vector for each protein sequence. The sequence-based feature vector of each gene pair was calculated by taking the absolute difference between the vectors of the proteins encoded by the two genes in the pair.

#### Protein-protein interactions

We retrieved protein-protein interactions (PPI) from the STRING database[37], considering only manually curated or experimentally validated interactions that were. We built a network graph of genes (nodes) and undirected interactions between them (edges), and extracted a 64-dimensional embedding vector for each gene (or node) in the network using the Node2Vec method with default parameters [38]. To obtain the PPI feature vector for each pair of genes, we took the absolute difference between the node embedding vectors of the two genes, similar to what was done for amino acid sequence.

#### Cancer cell line omics

We retrieved dependency scores of cancer cell lines measured upon gene perturbation from the Cancer Dependency Map portal (public release 2018q3 [27, 28]). Gene expression, mutation and copy number aberration data from the Cancer Cell Line Encyclopedia (CCLE) [29, 30] was obtained from the cBio portal repository [31, 32]. Based on these omics data, we defined alterations more broadly as encompassing non-silent mutations, gene expression z-scores larger than −1.96 or smaller than 1.96 (95% confidence), and discrete copy number aberration score equal to −2 (amplification) or 2 (deep loss). For gene expression, we used the log-transformed mRNA z-scores compared to the expression distribution of all samples (RNA-seq RPKM). For copy number scores, we used discrete values generated by the GISTIC algorithm [31, 32].

We created two feature sets based on cell line omics. The first one, *CRISPR with mutation*, was based on CRISPR gene dependency scores and mutation data. The second, *CRISPR with expression*, was based on CRISPR dependency scores and gene expression. Each of these feature sets comprised four features, namely the average dependency score of the first (or second) gene across cell lines where the second (or first) gene was unaltered, and average dependency score of the first (or second) gene across cell lines where the second (or first) gene was altered.

#### Tissue omics

We collected gene expression, mutation, copy number aberration, and clinical data for TCGA patient tissue samples [34] from the cBio portal[31, 32]. We used two different versions of gene expression scores: log-transformed mRNA z-scores compared to the expression distribution of all samples (RNA Seq RPKM) to identify alterations based on gene expression, and mRNA gene expression (RNA Seq V2 RSEM) to quantify gene expression level. Additionally, we collected healthy donor tissue gene expression data as transcript per million (TPM) from the GTEx portal [33] (dbGaP Accession phs000424.v8.p2).

Alterations were defined as encompassing non-silent somatic mutations, gene expression z-scores larger than 1.96 or smaller than −1.96 (95% confidence), and discrete copy number score equal to 2 (amplification) or −2 (deep loss). Using these alterations, we categorized patient tumour samples into two groups: samples with alterations in both genes, where alteration in one of the omics was sufficient; and the remaining samples without simultaneous alterations in both genes.

From tissue omics, we generated the following groups of features: patient survival, average gene expression in altered or unaltered tumour patient samples, gene coexpression in patient tumour/normal tissue or in healthy donor tissue, and correlation of copy number aberrations in patient tumour samples. The survival feature corresponded to the log-rank test p-value between the groups of patient samples with and without alterations in both genes, as defined above. Four average gene expression features were defined as the average gene expression of the first (or second) gene in tumour samples where the second (or first) gene was: unaltered (2 features) or altered (2 features). Additionally, six coexpression features were calculated as the Pearson’s correlation and respective p-value between the expression levels of the two genes in a gene pair in the following sets of samples: TCGA tumour samples from cancer patients (2 features), TCGA normal samples from cancer patients (2 features), and GTEx healthy donor tissue samples (2 features). Finally, two features expressing the correlation and p-value of copy number aberrations between the two genes in a gene pair was calculated using Spearman’s correlation.

#### Synthetic lethality labels

We obtained experimentally derived SL labels from the DiscoverSL [20], ISLE [13], Exp2SL [21], and Lu et al. [22] studies, which collectively aggregate the results of 25 original experimental studies (Supplementary Table S4). We removed all gene pairs for which there were any disagreements in SL label across the studies (Supplementary Table S1). Unless otherwise specified, we used one single SL dataset containing all unique gene pairs found across the four different sources of SL labels.

### ELISL models

ELISL models are mathematical functions that take as input the feature vector representation of a given gene pair, and generate an SL prediction score corresponding to the probability that such gene pair is synthetic lethal. The models are learnt using labelled gene pairs, with experimentally determined synthetic lethal status. In ELISL models, the representation of a gene pair comprises features from multiple, context-free and context-specific, data sources.

### Early late integration framework

The early late integrated framework is designed to learn models from a given number *k* of data sources, with *k* ∈ 𝕅 and *k* ≥ 2, as follows. We build *k* models, each learning from the feature set created for one of the *k* individual data sources of interest. Additionally, we train an additional model using the feature set obtained by concatenating the features generated from all the individual *k* data sources. The predictions of the *k* + 1 models are aggregated using weighted average based on the validation performances of the individual models. More formally, each individual dataset *X*_*i*_, with *i* ∈ 𝕅 and {1, …, *k*}, is a feature matrix 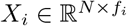, with *N* defined as the number of examples (rows in *X*_*i*_), *f*_*i*_ as the number of features (columns in *X*_*i*_). The concatenated dataset is defined as 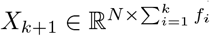 and results from concatenating the set of feature matrices for the *k* individual data sources, *X*_1_, …, *X*_*k*_. Each individual model is an ensemble of models learnt using a given dataset *X*_*i*_ with the corresponding labels for its *N* examples (gene pairs). Models are trained together with shared hyperparameters. Finally, the prediction score of a pair is calculated as 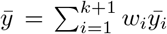, where *w*_*i*_ is the weight of model *i* and 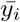 is the prediction probability score of the gene pair in model *i*. The weight *w*_*i*_ of each model in the final score is determined as the prediction performance on the validation set normalized over all models: 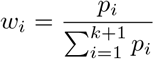, where *p*_*i*_ denotes the performance of model *i* (further details in Supplementary Information).

### Model training and evaluation

We learned ELISL models using two different types of ensembles of decision trees, random forests (ELISL-RF, [56]) and gradient-boosted decision trees (ELISL-GB, [57]). For each cancer type, we independently generated ten splits of the labelled pairs into disjoint train set (80%) and test set (20%). The pairs were drawn using random undersampling of the majority class to ensure a balanced number of examples or gene pairs with positive and negative SL labels.All SL prediction models, both ELISL and competing approaches, were evaluated in ten runs, each using one of the ten generated train/test set splits. First, the SL prediction models were learnt using the featurized representations and known SL labels for the gene pairs included in the train set. For ELISL models, the hyperparameters and the weight of each dataset were determined per run based on Bayesian grid search with 5-fold cross-validation, using AUPRC as measure of performance on the validation set (hyparameters in Supplementary Table S7). Finally, the SL prediction models learnt in every run were evaluated on the test set of that run using the AUPRC as performance measure.

### Systematic evaluation of all SL prediction models

#### Single cancer models and within cancer prediction

We trained and evaluated all SL prediction models within each cancer type, where train and test sets were constructed as usual such that they did not share gene pairs. As previously described, models were evaluated per cancer type over 10 runs using independently drawn train/test splits. This was performed for eight different cancer types: BRCA, CESC, COAD, KIRC, LAML, LUAD, OV, and SKCM. In addition to the ELISL models, ELISL-RF and ELISL-GB, we included five different methods recently proposed in the literature, which have shown high performances in different categories: matrix factorization approaches and graph-based methods, PCA-gCMF[15], GRSMF[16], and GCATSL[19]; and feature-based supervised learning approaches, SBSL-EN and SBSL-MUVR [14]. We tested for statistical significance of the difference in performance between the best ELISL model and the best of the other methods using the two-sided Wilcoxon signed-rank test.

### Robustness to gene selection bias

#### Double gene holdout (induced differences in train/test bias)

We again created 10 pairs of train and test sets, but this time ensured that there were no common genes between each train set and its corresponding test set. In other words, a gene could be in either the train or test set but not in both. This guarantees that train and test set necessarily follow different gene selection bias. We then evaluated all SL prediction methods on four cancer types: BRCA, CESC, LUAD, and OV. We excluded COAD, KIRC, LAML, and SKCM due to low performances, high variation in performances, and limited number of genes. For KIRC, the limited number of available gene pairs had already led to significant variation in performance across runs with the standard train/test splits allowing for individual gene overlap. For COAD and LAML, all SL prediction methods showed much lower performances than for the remaining cancer types. For SKCM, 99% of the pairs involved the MYC gene, thus leaving an insufficient number of genes to generate train/test sets without gene overlap. We tested for statistical significance of the difference in performance between the two methods that performed the best in the original experiment using standard train/test splits.

#### Cross-SL dataset prediction (inherent differences in train/test bias)

In this experiment, we trained the SL prediction models using SL labels from one source and tested them using SL labels from another source for the same cancer type. If a given gene pair appeared in both the train and test set, we removed this pair from the train set. For breast cancer (BRCA), we trained models on ISLE [13] labels and tested them on DiscoverSL [20] labelled pairs. For lung cancer (LUAD), we trained the model on DiscoverSL [20] labels and tested on labelled pairs from either Exp2SL [21] or Lu et al. [22]. We also conducted the versions of these experiments where we reversed the roles of the train and test set.

### Cross-cancer and pan-cancer prediction of ELISL-RF models

#### Cross-cancer prediction of single cancer models

We trained models per cancer type and evaluated their predictions separately on each of the other cancer types. For this, we used the same 10 pairs of train and test sets from the original single cancer experiment.

#### Pan-cancer model prediction

Pan-cancer models were obtained by ensembling the already trained models from each cancer type, where the weight of each model in the final prediction was attributed based on its validation performance. Combining the predictions of the different models allowed us to bypass simultaneous training using cancer types with largely different numbers of pairs. This would have required us to balance the data across cancer types, which could also severely limit the number of pairs available for training.

### Interpretation of the ELISL-RF models

#### Importance of feature sets

We calculated the importance of each feature set for the ELISL-RF models of the six cancer types with the least variance in AUPRC scores across runs. We considered the five feature sets used by the individual ensemble models that compose the ELISL-RF model, namely: sequence, PPI, cell line CRISPR with mutation, cell line CRISPR with expression, and tissue. To calculate the importance score for a given feature set, we permuted all of its features across the gene pairs in the test set, so as to break the relation between features and labels. When permuting a given feature set, the concatenated features also changed accordingly. We calculated the errors for the predictions in both the original test set and the permuted test set as (1-AUPRC) scores. The importance score was then defined as the ratio between the prediction error using the original test set and the prediction error using the permuted test set.

### Impact of high-dimensionality of sequence embeddings

To analyze the impact of the high-dimensionality of the sequence feature representation, we re-trained the single-cancer ELISL-RF models using different sequence embedding sizes and re-evaluated the performance. We evaluated the following embedding sizes: 32, 64, 128, 256, 512, and 1024.

### Analysis of newly predicted SL gene pairs

In addition to evaluating the prediction performance for gene pairs with known labels (test set), we assessed the potential of ELISL-RF to generate promising new SL gene pairs. We did this by generating a set of gene pairs with no known labels and then predicting their probability of being synthetic lethal. We used the breast cancer ELISL-RF model, since it was one of the best performing methods across experiments with similar and different gene selection bias between train and test set (Fig. 1 and Fig. 2).

#### Generating a set of unknown gene pairs

We created the set of unknown pairs by generating all pairwise combinations between genes found in cancer and DNA repair pathways. We retrieved KEGG, PID, and Reactome pathway gene sets from the molecular signatures database v7.1 [58, 59]. From the total of 572 genes found across all pathways (Supplementary Table S10), we generated 163, 306 gene pairs. After excluding the pairs already present in the train or test sets, we ended up with 163, 118 gene pairs.

#### Survival analysis of newly predicted SL gene pairs

To validate predicted SL gene pairs without known labels, we investigated if there were differences in survival time between patients with or without a simultaneous mutation in both genes of the unknown pairs predicted by ELISL-RF. We found that typically only a small number of patient tumour samples from TCGA carried simultaneous mutations in both genes, since tumour cells are not likely to survive when the gene pair is synthetic lethal. The sample sizes were too small to test for survival differences, so we decided to look at gene families rather than single genes. We stratified the patient tumour samples into two groups based on simultaneous mutation status, that is, presence or absence of deleterious mutations simultaneously in genes from both families. Specifically, for a given pair of genes (Gene 1, Gene 2), we denote the group of samples with mutations in both a member from the family of Gene 1 and a member from the family of Gene 2 as (Gene 1 and Gene 2), while the group without simultaneous mutations is expressed by ∼ (Gene 1 and Gene 2). Survival times of both groups were estimated using a Cox proportional hazards model, including covariates for age, sex, and cancer type in addition to simultaneous mutation status. We generated plots with the Kaplan-Meier survival curves for the two groups. For completeness, we also represented two subgroups of the group without simultaneous mutations, namely: the subgroup with a deleterious mutation in only one of the families but not both (Gene1 xor Gene2), and the subgroup with no deleterious mutation in any of the genes from both families (Unaltered). Note that, although the ELISL-RF model included a survival-based feature as part of the tissue-specific model, the contribution of the tissue features to the final ELISL-RF model was reportedly negligible. One reason for this is that the survival feature was very sparse due to the already mentioned rare occurrence of simultaneous mutations in both genes.

## Notes

### Competing Interest Statement

The authors have declared no competing interest.

