## Supplementary Information for "ELISL: Early-Late Integrated Synthetic Lethality Prediction in Cancer"

### ELISL Supplementary Materials

September 19, 2022

**4 Supplementary Tables** **23**

**5 References** **29**

### 1 Supplementary Results

#### 1.1 Impact of Sequence Dimension

Besides the training labels and algorithm itself, the high-dimensional feature space can also cause bias by allowing the model to overfit [2]. Since the sequence dataset we generate is 1024-dimensional, we investigated if the high dimension impacts the performance. We repeated the single cancer experiment with ELISL-RF by reducing the dimension of the sequence dataset each time. We also expected a drop in the performance due to linear transformation from PCA; however, except for ovarian cancer, the performance changes were within the standard deviation of the original experiment Supplementary Figure S6. The mean performance on ovarian cancer dropped from 0.87 AUPRC scores to 0.8 AUPRC scores from dimension-1024 to dimension-32. We already know that ovarian cancer is prone to bias; however, the performance was still the same on dimension-512 and dimension-256. Additionally, the performance of ovarian did not drop drastically, and the drop was acceptable. Therefore, the high-dimension of sequence dataset was not the reason behind our high performance.

#### 1.2 Extra Survival Analysis

Before analyzing the survival of patients when there is simultaneous mutation in the predicted pairs, we first analyzed a well-known synthetic lethal pair in breast cancer: BRCA1-2 and PARP1-2(Supplementary Figure S9). Due to the limited number of samples with a simultaneous mutation in both groups, we analyzed the PARP family(1 to 16) instead of only looking at PARP1-2. Patients with simultaneous mutation in both of the BRCA(1-2) group or the PARP(1 to 16) family have 20 months more median survival time than the rest of the patients(Supplementary Figure S9). Additionally, to confirm this is not the case for the pairs with very low scores, we also analyzed the survival of 4 pairs that have the least score (Supplementary Figure S8). PARP1|RIPK1 and MAP3K7|PARP1 are the negatively labeled pairs with the least scores. For both of the pairs, the survival of the patients who have a simultaneous mutation in the family(PARP, RIPK, MAP3K families) of both genes have worse survival than the rest of the patients (Supplementary Figure S10-S11). Moreover, we investigated the least scored pairs without a label: MAP2K2|PARP1 and DAPK2|PARP1 (Supplementary Figure S12-S13). While the interaction was the same between DAPK and PARP families, patients with simultaneous mutation in both MAP2K and PARP families had better survival. However, since these unknown pairs don't have any experimental labels, they are not confident as negatively labeled pairs. By inferring these results on the well-known SL pair, we expect the top-scored unknown pairs to have the same characteristics as the BRCA-PARP pair.

### 2 Supplementary Methods

#### 2.1 Dataset Generation Details

##### Cell line datasets

We first defined alterations from different omic data:

- **mut**: If a gene is mutated there is alteration excluding silent mutations from mutation omic.

- **expr**: All RPKM gene expressions values are converted to Z-score. If z-score of gene is not between (-1.96, 1.96), then altered.
- **cnv**: Copy number alteration discretized by GISTIC. Alteration if cnv>2 or cnv<-2

We then defined retrieve dependency scores for each gene cell line pair from the sources: crispr or d2(RNAi) dependency scores.

We generated five different types of datasets from cell line data:

- **source_dep_omic**: source as dependency matrix, omic as alteration source
  - **dep0**: Avg. dependency score of gene1 when gene2 altered
  - **dep1**: Avg. dependency score of gene1 when gene2 not altered
  - **dep2**: Avg. dependency score of gene2 when gene1 altered
  - **dep3**: Avg. dependency score of gene2 when gene1 not altered
- **source_dep_any**: source as dependency matrix, alteration source is all omics together.
  - **dep0**: Avg. dependency of gene1 when gene2 altered in any omic
  - **dep1**: Avg. dependency of gene1 when gene2 not altered in any omic
  - **dep2**: Avg. dependency of gene2 when gene1 altered in any omic
  - **dep3**: Avg. dependency of gene2 when gene1 not altered in any omic
- **source_dep_muex**: source as dependency matrix, alteration source is mutation and gene expression omics.
  - **dep0**: Avg. dependency of gene1 when gene2 altered
  - **dep1**: Avg. dependency of gene1 when gene2 not altered
  - **dep2**: Avg. dependency of gene2 when gene1 altered
  - **dep3**: Avg. dependency of gene2 when gene1 not altered
- **source_dep_comb**: Concatenation of datasets source_dep_omic where omic is mut, expr and cnv.
- **comb_dep_x**: Concatenation of **crispr** and **d2** datasets with alteration as x.

The list of datasets we created using these types:

- **source_dep_omic**:
  - crispr_dep_mut
  - crispr_dep_expr
  - crispr_dep_cnv
  - d2_dep_mut
  - d2_dep_expr
  - d2_dep_cnv
- **source_dep_any**:
  - crispr_dep_any
  - d2_dep_any
- **source_dep_muex**:
  - crispr_dep_muex
  - d2_dep_muex
- **source_dep_comb**:
  - crispr_dep_comb
  - d2_dep_comb
- **comb_dep_x**:
  - comb_dep_mut
  - comb_dep_expr
  - comb_dep_cnv

- comb_dep_any
- comb_dep_muex
- comb_dep_comb

We then evaluated these datasets in our training set and decided to use only crispr_dep_mut, crispr_dep_expr as cell line dataset for our applications.

###### Tissue dataset

For the tissue, we only defined one dataset. To generate some of the features for the tissue dataset, we defined alterations from different omic data similar to cell lines:

- **mut**: If a gene is mutated, there is alteration excluding silent mutations from mutation omic.
- **expr**: All RPKM gene expressions values are converted to Z-score, and if the z-score of a gene is not between (-1.96, 1.96), then it is altered.
- **cnv**: Copy number alteration discretized by GISTIC. There is alteration if cnv>2 or cnv<-2.

For a detailed description of how z-scores are computed and copy number changes are discretized, refer to cBioPortal File Formats.

The features of the tissue dataset are:

- **surv**: Log-rank test p-value between two groups: Patients who have simultaneous alteration in both of the genes in pair. Alteration in at least one omic is enough.
- **expr0**: Avg. gene expression of gene1 when gene2 is not mutated.
- **expr1**: Avg. gene expression of gene1 when gene2 is mutated.
- **expr2**: Avg. gene expression of gene2 when gene1 is not mutated.
- **expr3**: Avg. gene expression of gene2 when gene1 is mutated.
- **t_coexp**: Coexpression of two genes between tumor tissues (Pearson correlation).
- **t_coexp_p**: p-value of coexpression of two genes between tumor tissues (Pearson correlation).
- **t_cocnv**: Cocnv of two genes between tumor tissues (Spearman correlation with discretized copy number alteration).
- **t_cocnv_p**: p-value of cocnv of two genes between tumor tissues (Spearman correlation with discretized copy number alteration).
- **n_coexp**: Coexpression of two genes between normal tissues (Pearson correlation).
- **n_coexp_p**: p-value of coexpression of two genes between normal tissues (Pearson correlation).
- **h_coexp**: Coexpression of two genes between healthy tissues (Pearson correlation).
- **h_coexp_p**: p-value of coexpression of two genes between healthy tissues (Pearson correlation).

#### 2.2 Model Details

##### 2.2.1 Dataset Contribution Weights

Final prediction score of a pair is $\bar{y} = \sum_{i=1}^{k+1} \frac{w_i}{w_t} \bar{y}_i$ where $w_i$ is the weight of model $i$, $w_t = \sum_{i=1}^{k+1} w_i$, $\bar{y}_i$ is the prediction probability score of the pair in model $i$ and $k$ is the number of dataset. The concatenated dataset is the $(k+1)$th dataset. We define the weight of model

$i(w_i)$ as the AUPRC score of the model on validation set if there is. In the cases where we don't have validation set we use AUPRC score on the training set itself.

While investigating dataset combinations using only the training set, we applied 5 fold cross-validation to the training set and did not tune the parameters. Since we also divided the dataset into 20% test and 80% at each fold during these experiments, we did not have enough samples for validation. Therefore we only considered the training AUPRC score as the weight for each model.

For the other experiments, since we use the whole training set for training, we were able to do grid search 10-fold cross-validation, and we used the mean AUPRC of the 10-fold cross-validation as model weights. Since we only use the training set for these, we tuned the parameters and found the weights of models without looking at the test set. Additionally, while calculating the scores, we normalized the weights by dividing the weight by the total weight to predict the probability between 0 and 1.

###### 2.2.2 Evaluation

To evaluate our model, we did not use a traditional accuracy score since it can have an unreliable and inconsistent result for some cases like unbalanced data[9]. Therefore, we used more reliable evaluation methods Area under the precision-recall curve(AUPRC), the area under the receiver operating characteristic curve (AUROC), which are also not dependent on the cutoff and works with prediction probability directly. While ROC visualizes true-positive rate(TPR) versus false-positive rate(FPR), PRC visualizes precision versus recall for all possible cutoff values. Then both of the methods calculate the area under curves(AUC). The difference between AUROC and AUPRC is that AUPRC does not take correctly predicted negative pairs into account. If a problem aims to identify the positive samples more than the negatives, then AUPRC is a better choice. Therefore, although we evaluate our methods with both AUPRC and AUROC, we use AUPRC as the primary evaluation score since, for our problem, the main goal is to find positive synthetic lethality pairs. Both of the scores are between 0 and 1, but while random performance is always 0.5 for AUROC, it is the ratio of positive samples over all samples for AUPRC. Therefore, in extreme cases, it is hard to evaluate AUPRC correctly. Table S2 shows the ratio of positive samples for each cancer type. For some cancer types, the random performance is less than 0.01, which is hard to interpret. Thus, we also balance the test dataset during our experiments since negative pairs are not essential and reduce the AUPRC score's interpretability.

We also investigate our results with a recent performance metric, Matthews Correlation coefficient(MCC) score, which needs binary predictions. Due to unreliable scores of accuracy on problems with the class imbalance and asymmetric nature of other binarized metrics like precision, recall, and F1-score, MCC is proposed to deal with these problems and firstly used in secondary structure prediction[14]. It calculates the correlation between prediction and the true class labels and has the same properties as the Pearson correlation coefficient. Thus, it changes between -1 and 1, where -1 is the reverse correlation, 0 is no correlation, and 1 is high-correlation[1]. We evaluated our methods by choosing the cutoff as 0.5 since it indicates the class with a higher probability score since it is binary classification.

###### 2.2.3 Optimization

We finetune the parameters at each run separately. For each run, we tune the parameters by applying Bayesian Optimization with Gaussian Process to minimize the error(1-AUPRC) on the validation set. We find the error score on 10-fold cross-validation on the training set of each run. Additionally, we did not finetune each Random Forest model separately. Thus, the parameters are shared across all Random Forest (or Gradient Boosting Decision Tree) models for each run. The parameter details can be found in (Table S7)

###### 2.2.4 Importance of Datasets

To calculate the importance of each dataset, we used a technique similar to permutation feature importance. By permuting the rows of feature matrix we disconnect labels connection with all of the features in the dataset. The detailed algorithm is given in Algorithm 1:

**Algorithm 1** Permutation Dataset Importance

**Require:** Trained model $m$, set of feature matrices $X_i$ from each dataset $i$ for test.
**Ensure:** $score\_list_i = list()$

**for** `each fold` **do**
  $error \leftarrow 1-$AUPRC(original datasets)
  **for** `each matrix` $X_i$ `from dataset` $i$ **do**
    **for** `20 times` **do**
      `Permute the dataset` $X_i$
      $pert_error_i \leftarrow 1-$AUPRC(only $X_i$ is permuted)
      $score\_list_i$`.append(`$pert\_error_i/error$`)`
    **end for**
  **end for**
**end for**
**for** `each matrix` $X_i$ `from dataset` $i$ **do**
  $mean_i$, $std_i \leftarrow score\_list_i$
**end for**

###### 2.2.5 Tools

All of the codes for experiments are written and integrated with Python3.6. Only PCA-gCMF and Seale methods were run in R[17].

We used **LightGBM**[12] and **scikit-learn**[16] together for the Regularized Random Forest and Regularized Gradient Boosting Decision Trees.

We optimize the model using the Bayesian Optimization with Gaussian Process in **scikit-optimize**[7] package.

For the visuals, we made use of **seaborn**[19] and **matplotlib**[10] libraries. Additionally, for the Kaplan-Meier graphs and tests regarding survival, we used **lifelines**[3] package.

##### 2.3 Other Methods

While we also use R for PCA-gCMF and Seale method, the other methods are integrated into our framework using Python.

###### PCA-gCMF

PCA-gCMF is a version of collective matrix factorization(CMF). It uses principle component analysis(PCA) to deal with identical rows and columns across multiple matrices and group-sparse CMF to decompose these multiple matrices. We only chose PCA-gCMF to compare since it was the best performing method in their work. They also have different methods such as CMF, PCA-CMF, gCMF, and PCA-gCMF. They use a gene dependency profiles matrix for gene pairs, where they use the p-value of the Wilcoxon rank-sum test to find the change in dependency score of a gene when there is a mutation or not in the second gene in the pair. They also use mRNA expression count matrix (genes $\times$ samples), Co-expression matrix (genes $\times$ samples), which measures the Spearman correlation coefficient between the expression of two genes in all cohort and lastly, the CNV profile matrix (genes $\times$ samples) where the values are the continuous copy number variance.

###### GRSMF

Graph regularized self-representative matrix factorization(GRSMF) is another matrix factorization method that makes use of GO biological process(BP) similarity matrix as a regularizer[8]. We constructed the similarity matrix as described in their paper. Although they select the optimal values for their parameter using grid search, this search is done on a test set which maximizes the performance. Since we define the parameters before seeing the testing performance, we did not tune their parameter and use the parameters mentioned in the paper.

###### GCATSL

GCATSL is one of the first methods to use synthetic lethal labels to create a network and find missing interactions[13]. They use contextualized(featured) attention networks, diffuse features into local-global neighbors, and train neural networks to construct, predict the SL network. They use 3 different similarity matrices as datasets: GO Biological Process similarity, GO Cellular Component similarity, and PPI similarity. In their work, they use BioGrid[15] as a PPI graph source, but we used STRING[11] as a PPI source to make the comparison fair since we use STRING as a PPI network source. We used their default hyperparameters since, for their work, they did not search for the best hyperparameters blindly(Without looking at test performance).

###### Seale Methods

Seale methods includes 2 linear(Regularized Random Forest(RRF)[4], Multivariate methods with Unbiased Variable(MUVR)[18]) and 2 non-linear models (L0L2[6], Elastic Net[5]) with same set of features[17]. They created one dataset from hand-crafted features using tissue and cell line omics and also pathway gene sets. They additionally investigated the biases in these synthetic lethality experiments. They applied grid search for all of their models to tune hyperparameters. We also applied the same techniques mentioned in their documents for tuning. Their features also include features from previous works, and we accept Seale models as the newest model with feature-based techniques.

##### 2.4 Links of Sources Used

###### Tissue Data

TCGA tissue omics and clinical data are taken from TCGA Firehose-cBioPortal
GTEX healthy tissue gene expression is taken from GTEx Portal Gene TPMs

GTEX annotation is taken from A de-identified, open access version of the sample annotations available in dbGaP.

##### Cell Line Data

Cell line omics are taken from CCLE_BROAD_2019
Crispr dependency scores are taken from Crispr Dependencies
RNAi(Demeter2) dependency scores are taken from RNAi Dependencies

##### PPI and Sequence Data

PPI graph is downloaded from STRING
Node to vector conversion is done with Node2Vec
Human proteins with reviewed amino acid sequence are downloaded from Uniprot
Sequence to vector conversion is done with SeqVec

### 3 Supplementary Figures

#### 3.1 Labels for Each Cancer Type

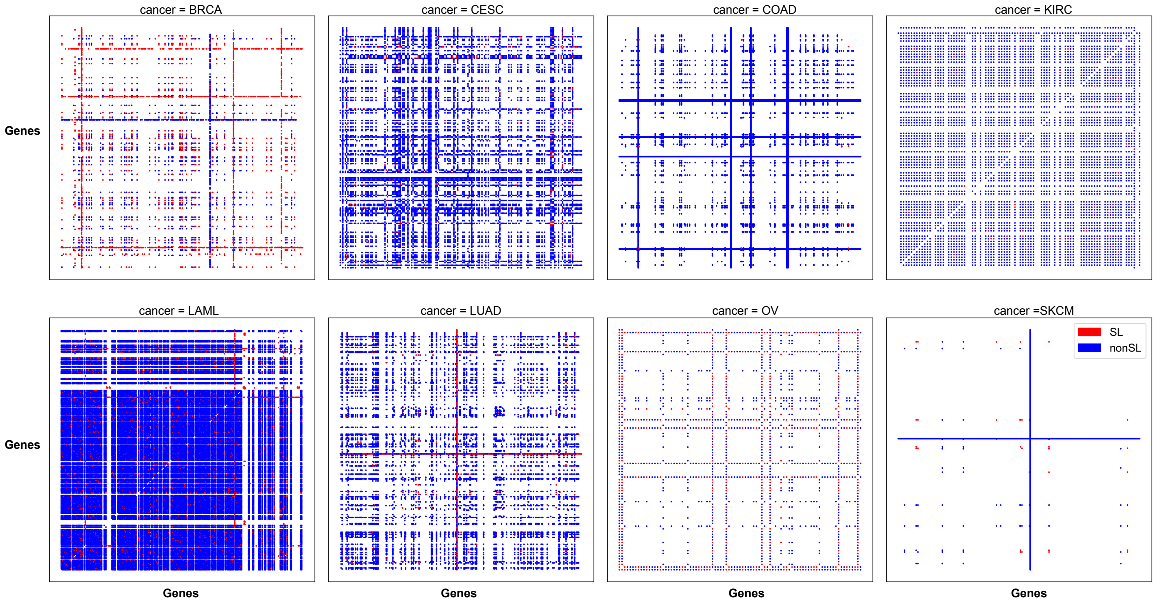

Figure S1: **Label distribution for each cancer type.** The positive(SL) and negative(nonSL) label distributions show unbalanced nature in these label sources. While some genes are investigated more, some of them are less investigated. Additionally, the number of genes investigated and how dense the labels are different for each cancer type. While LAML is the most dense, SKCM is the most sparse dataset. KIRC and OV are the cancer types with least number of different genes.

##### 3.2 Cross Validation vs. Training AUPRC Score

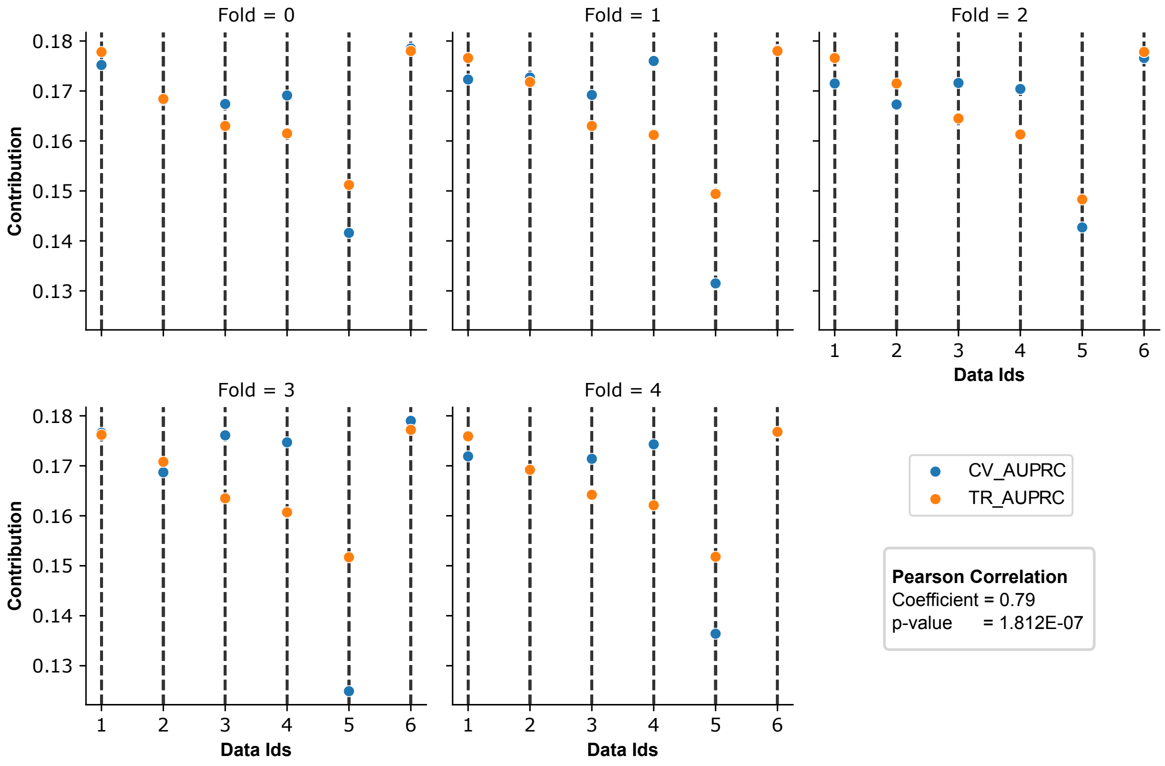

Figure S2: **Comparison of contribution techniques.** For the 5-fold cross-validation within the training set, we check both the AUPRC score on training and the validation set at each fold. We investigated if there is a drastic change in using training and validation performance while calculating contributions. For most of the cases, there is only a little change in contributions. Additionally, there is a high correlation between both of the contribution types(Pearson correlation coefficient=0.79). Data Ids refers to 1-Sequence, 2-PPI, 3-Crispr dependency with mutation, 4-Crispr dependency with expression, 5-Tissue, 6-Concatenated dataset.

#### 3.3 Single Cancer Experiment Extra Evaluation

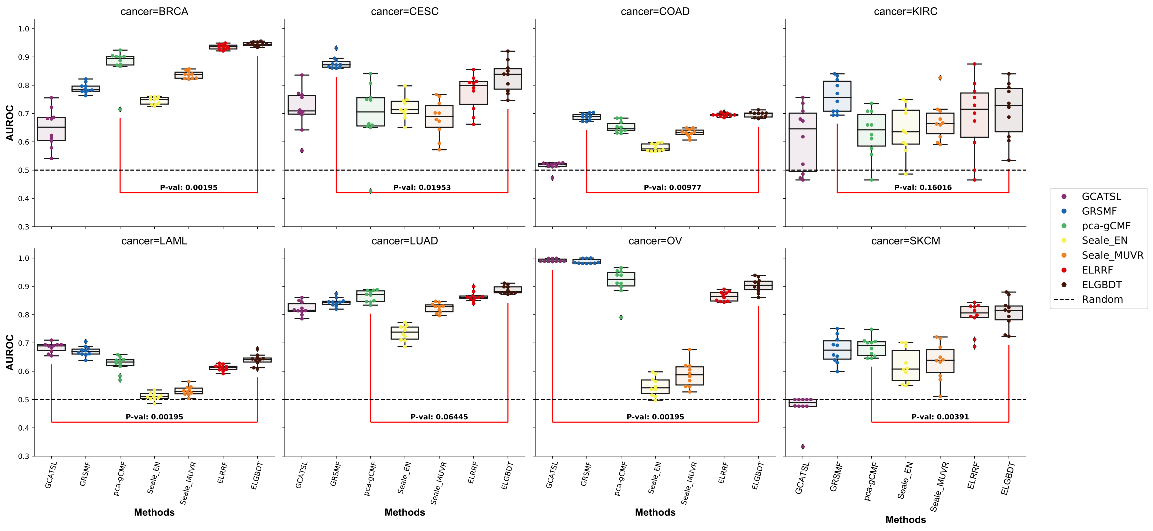

(a)

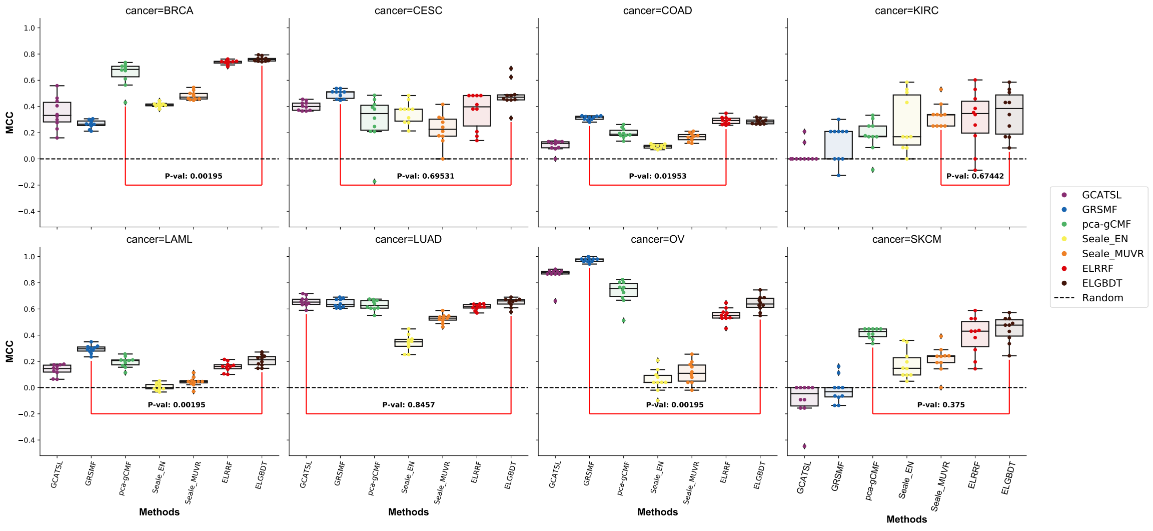

(b)

Figure S3: **Single cancer experiment results with other scores.** The p-value belongs to Wilcoxon signed rank test between the most successful ELISL method and the most successful of other methods. **a,** Results with AUROC performance. Similar to AUPRC results but our method is the best performer in 4 cancer types, which was 3 with AUPRC. **b,** Results with MCC performance which is again similar to results with other scores. Our method is the best performer in 4 cancer types and competitive in others.

#### 3.4 Double Holdout Experiment Extra Evaluation

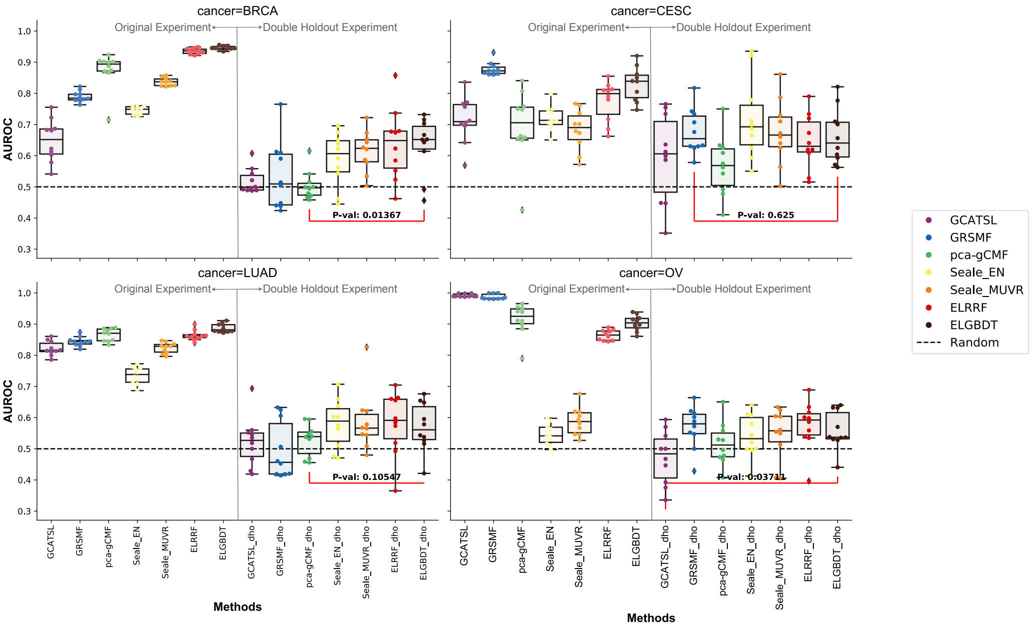

(a)

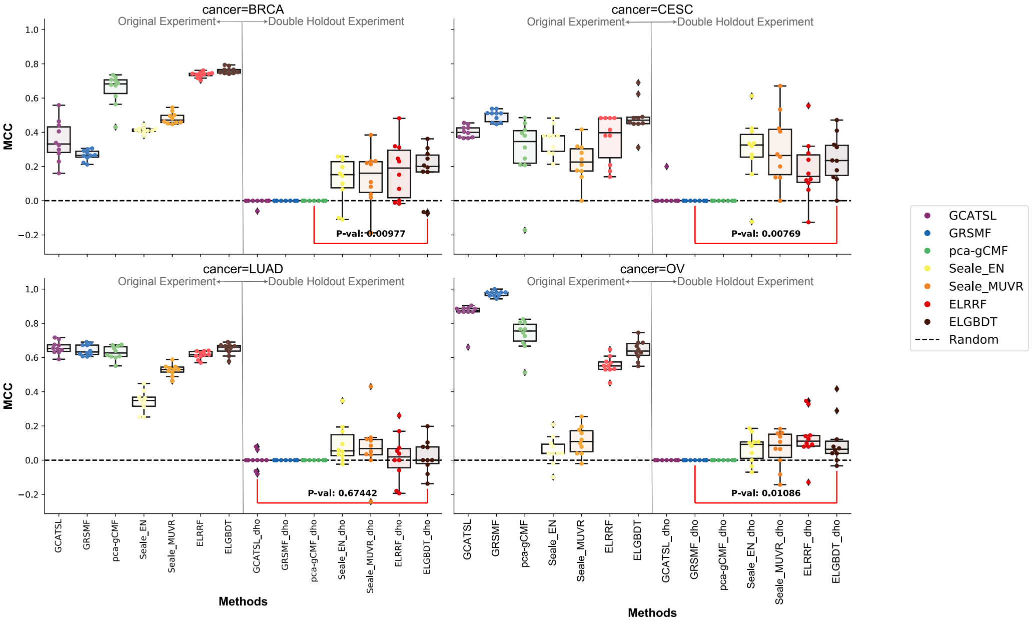

(b)

Figure S4: **Double holdout cancer experiment results with AUROC(a) and MCC(b) performance.** Similar to AUPRC results, our methods perform better than random for each case where matrix factorization or network-based models fails to perform better than random. The p-value belongs to the Wilcoxon signed-rank test between the most successful ELISL method and the most successful of other methods from the single cancer experiment.

##### 3.5 Cross Dataset Experiment AUROC Evaluation

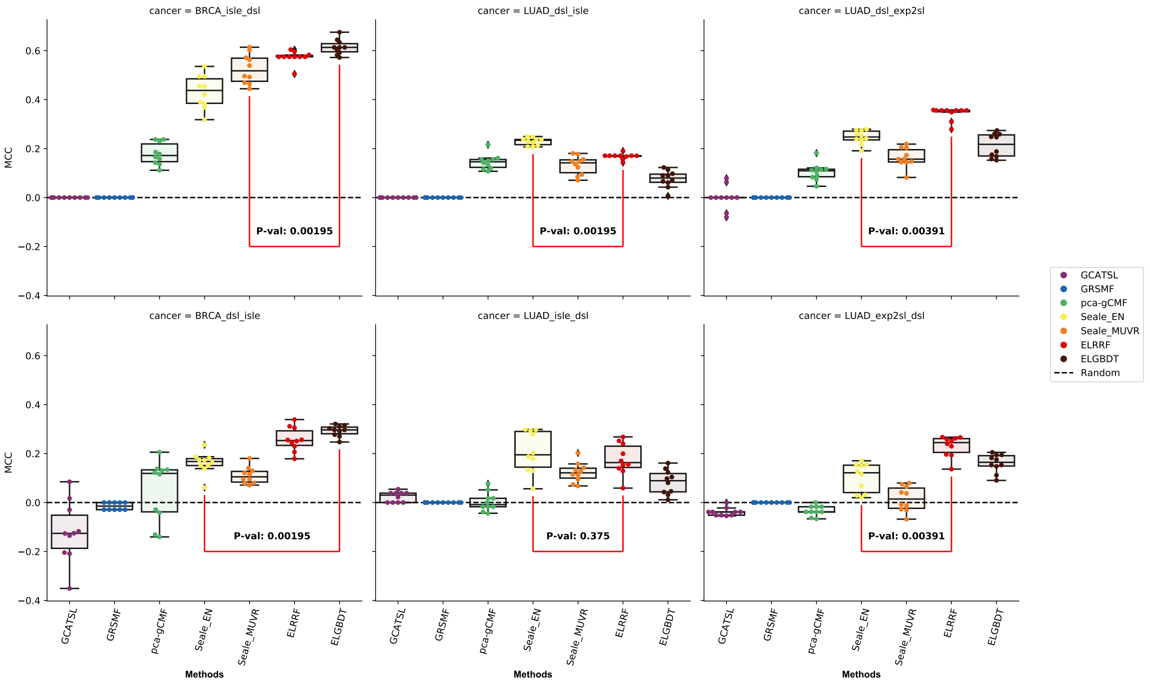

Figure S5: **Cross dataset experiment results with AUROC score.** The p-value belongs to Wilcoxon signed rank test between the most successful ELISL method and the most successful of other methods.

##### 3.6 Impact of Sequence Dimension

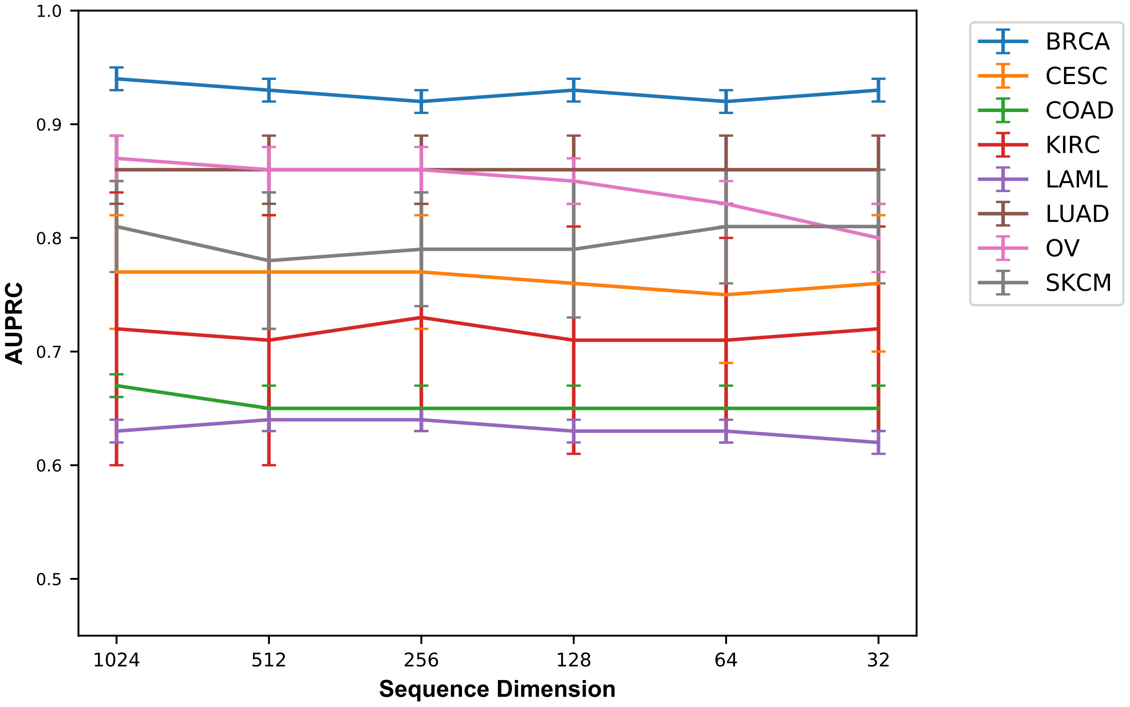

Figure S6: **Impact of sequence dimension to model performance** AUPRC Performance of ELISL-RF model as the dimension of sequence dataset is reduced from 1024 to 32. The vertical lines are the standard deviation of 10 runs. Reducing the dimensions of sequence dataset did not cause a drastic drop in performance.

##### 3.7 Cross Cancer Experiment AUROC Evaluation

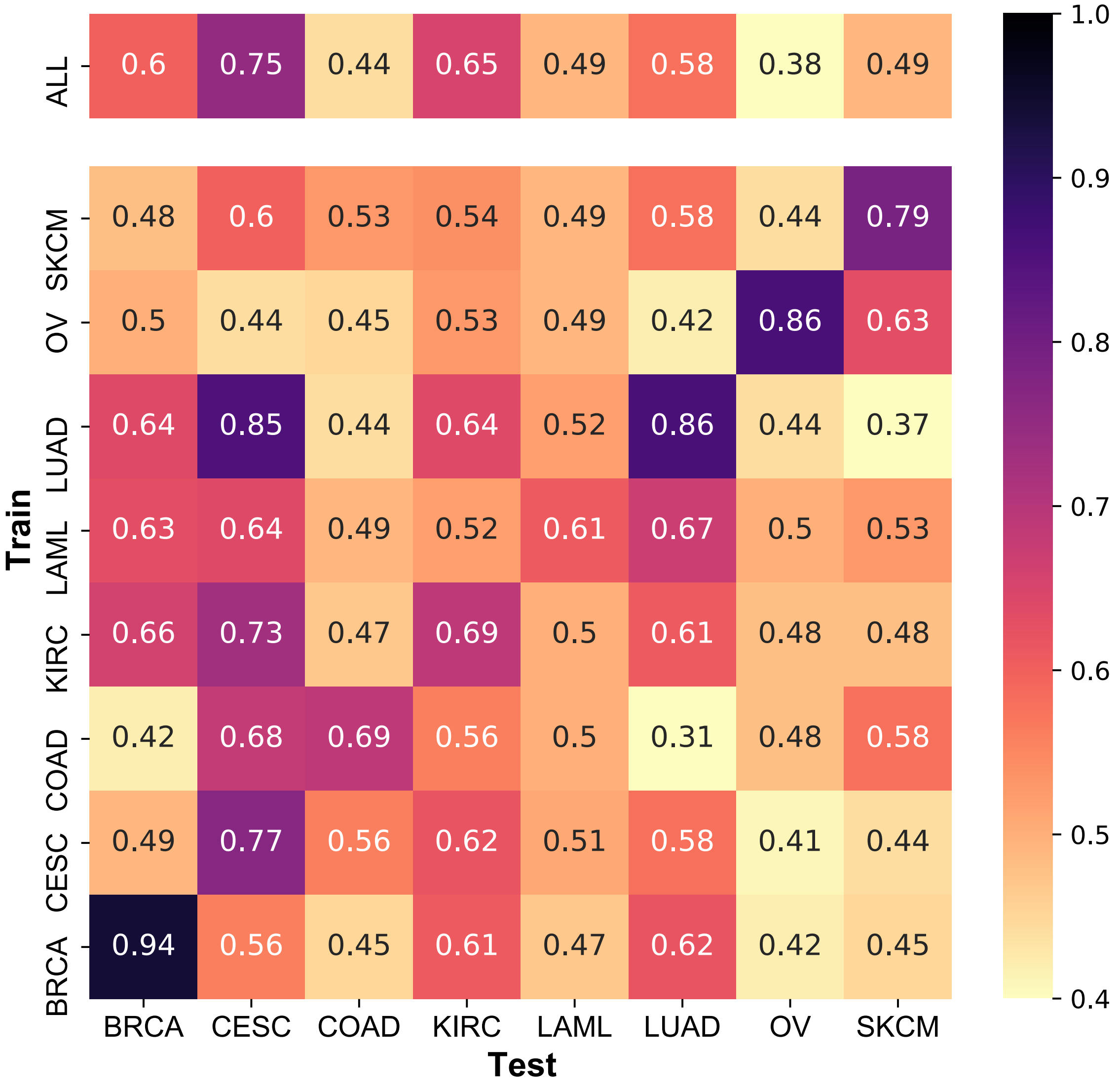

Figure S7: **Cross cancer and ensemble-cross cancer experiment results with mean AUROC score.** The top row is the AUPRC performance of predictions for each cancer type by ensemble of trained models of other cancer types.

#### 3.8 Bottom 10 Predictions with Testing and Unknown Pairs

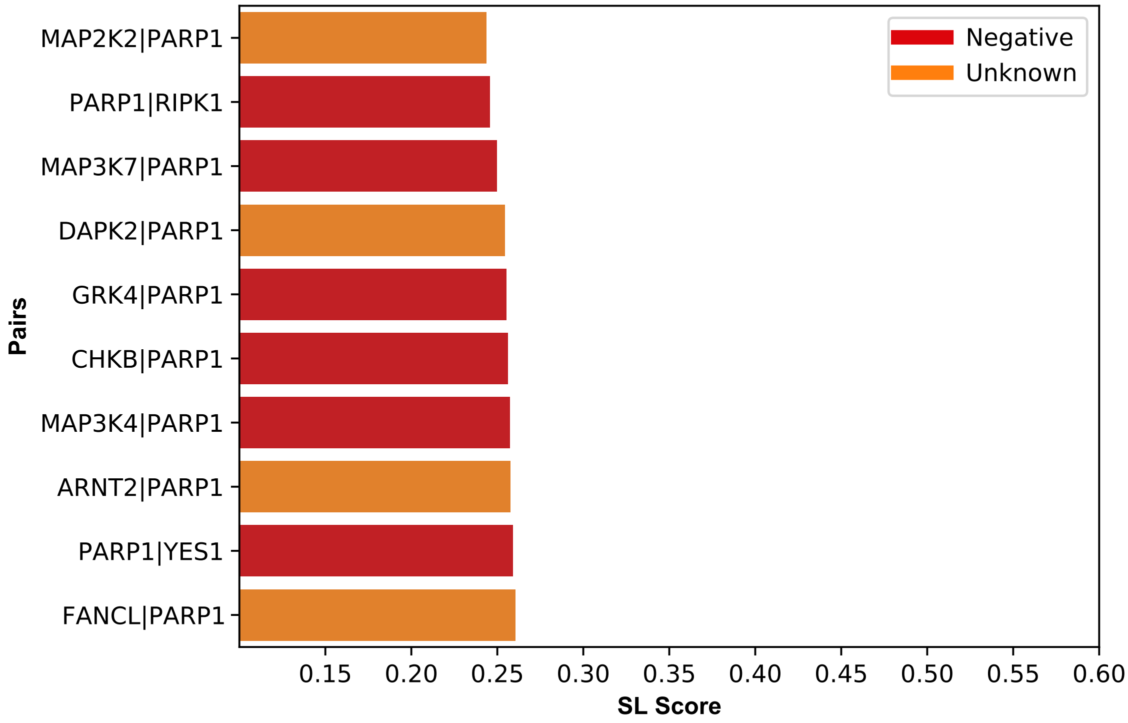

Figure S8: **Bottom 10 pairs in unknown experiments.** Least scored 10 pairs (Labeled in test set+Unknown) in single cancer experiment for BRCA with ELISL-RF.

#### 3.9 Survival Curves with Simultaneous Mutation

##### 3.9.1 BRCA-PARP Survival Analysis on Simultaneous Mutation

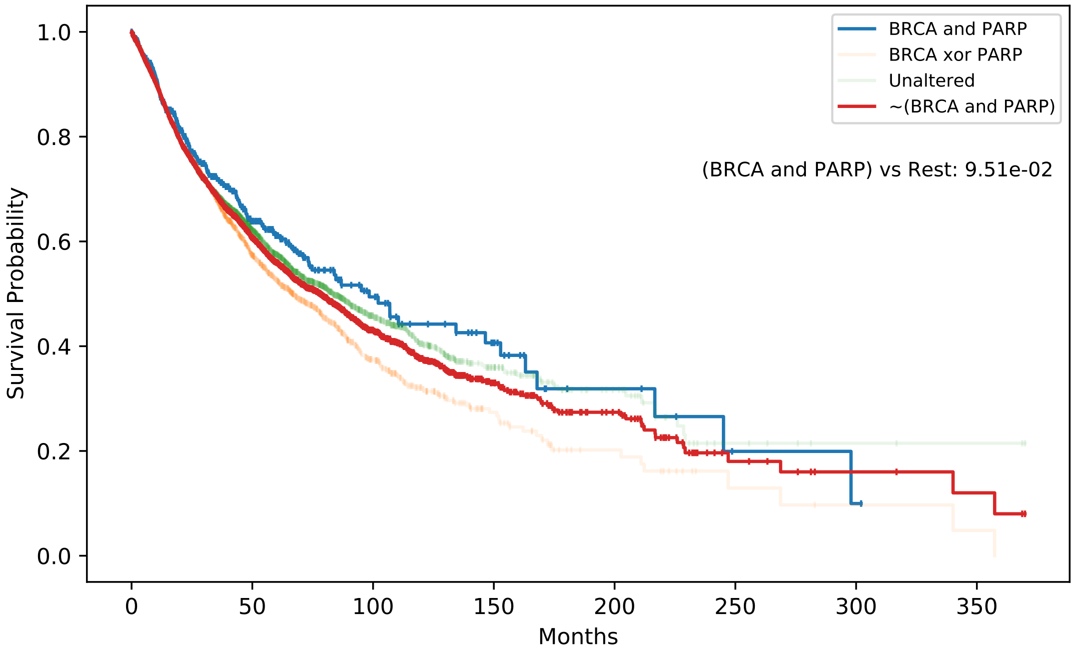

| | # of Cases | # of Events | Median Survival Time |
|---|---|---|---|
| BRCA and PARP | 501 | 160 | 98.4 |
| BRCA xor PARP | 3092 | 1102 | 66.67 |
| Unaltered | 7226 | 2263 | 83.31 |
| ~(BRCA and PARP) | 10318 | 3365 | 78.21 |

| | coef | exp(coef) | z | p |
|---|---|---|---|---|
| Diagnosis Age | 0.01 | 1.01 | 15.02 | 5.06e-51 |
| Cancer Type | -0.01 | 0.99 | -5.03 | 5.01e-07 |
| Mutation | 0.15 | 1.16 | 2.95 | 3.15e-03 |
| Gender | 0.09 | 1.09 | 4.31 | 1.66e-05 |

Figure S9: **Survival Analysis of simultaneous mutation in BRCA and PARP groups.** Kaplan Meier survival curve for BRCA(1-2) and PARP(1 to 16) and log-rank test p-value between the groups: Simultaneous mutation between two groups and the rest of the patients. The median survival time of the group with simultaneous mutation is 20 months more than the rest of the patients. The table at the bottom shows the coxph hazard function where *mutation* parameter is simultaneous mutation between two groups.

##### 3.9.2 RIPK-PARP Survival Analysis on Simultaneous Mutation

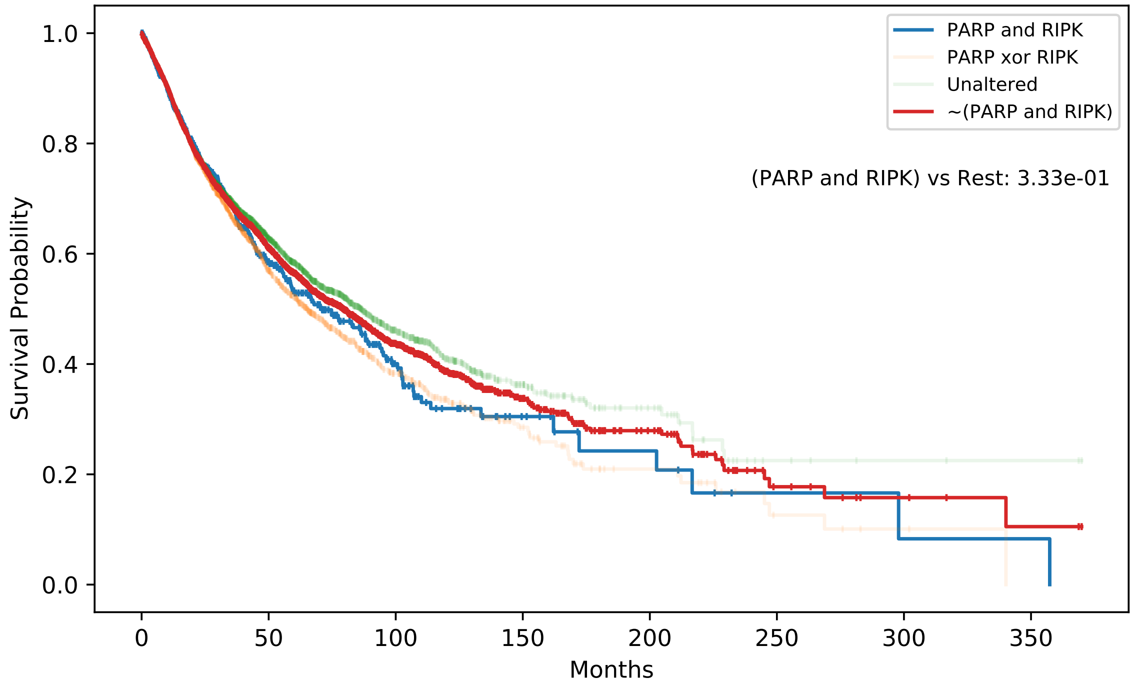

| | # of Cases | # of Events | Median Survival Time |
|---|---|---|---|
| PARP and RIPK | 836 | 296 | 70.82 |
| PARP xor RIPK | 2847 | 1025 | 65.0 |
| Unaltered | 7136 | 2204 | 86.27 |
| ~(PARP and RIPK) | 9983 | 3229 | 79.46 |

| | coef | exp(coef) | z | p |
|---|---|---|---|---|
| Diagnosis Age | 0.01 | 1.01 | 14.91 | 2.76e-50 |
| Cancer Type | -0.01 | 0.99 | -5.16 | 2.41e-07 |
| Mutation | 0.03 | 1.03 | 0.87 | 3.83e-01 |
| Gender | 0.09 | 1.09 | 4.34 | 1.43e-05 |

Figure S10: **Survival Analysis of simultaneous mutation in RIPK and PARP groups.**Kaplan Meier survival curve for RIPK(1 to 16) family and PARP(1 to 16) and log-rank test p-value between the groups: Simultaneous mutation between two groups and the rest of the patients. The median survival time of the group with simultaneous mutation is 9 months less than the rest of the patients. The table at the bottom shows the coxph hazard function where *mutation* parameter is simultaneous mutation between two groups.

##### 3.9.3 MAP3K-PARP Survival Analysis on Simultaneous Mutation

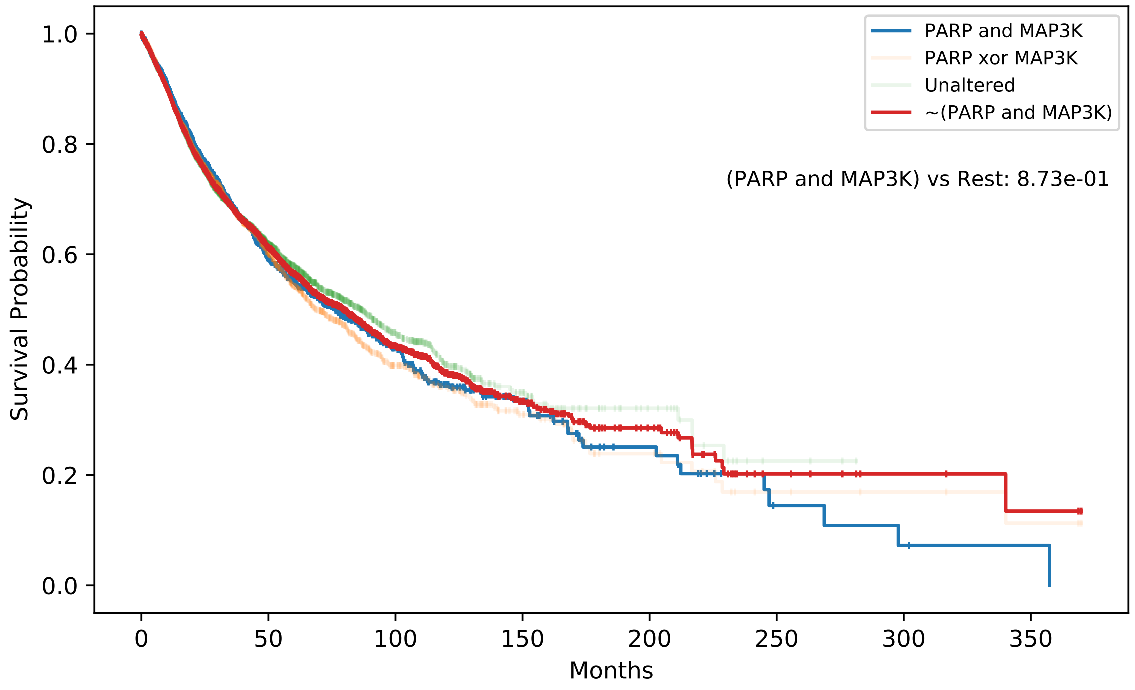

| | # of Cases | # of Events | Median Survival Time |
|---|---|---|---|
| PARP and MAP3K | 1823 | 645 | 76.24 |
| PARP xor MAP3K | 3422 | 1130 | 69.07 |
| Unaltered | 5574 | 1750 | 86.76 |
| ~(PARP and MAP3K) | 8996 | 2880 | 79.53 |

| | coef | exp(coef) | z | p |
|---|---|---|---|---|
| Diagnosis Age | 0.01 | 1.01 | 15.16 | 6.64e-52 |
| Cancer Type | -0.01 | 0.99 | -4.95 | 7.41e-07 |
| Mutation | 0.12 | 1.13 | 4.61 | 4.07e-06 |
| Gender | 0.09 | 1.09 | 4.30 | 1.67e-05 |

Figure S11: **Survival Analysis of simultaneous mutation in PARP and MAP3K groups.** Kaplan Meier survival curve for PARP(1 to 16) and MAP3K(1 to 16) family and log-rank test p-value between the groups: Simultaneous mutation between two groups and the rest of the patients. The median survival time of the group with simultaneous mutation is 3 months less than the rest of the patients. The table at the bottom shows the coxph hazard function where *mutation* parameter is simultaneous mutation between two groups.

##### 3.9.4 MAP2K-PARP Survival Analysis on Simultaneous Mutation

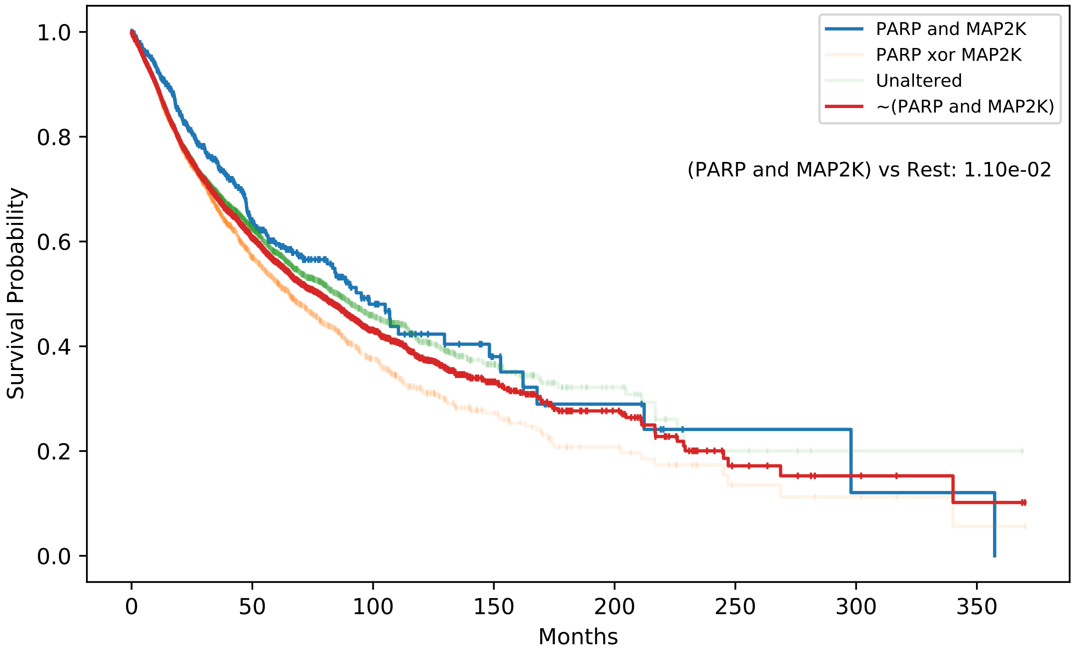

| | # of Cases | # of Events | Median Survival Time |
|---|---|---|---|
| PARP and MAP2K | 576 | 174 | 95.14 |
| PARP xor MAP2K | 3218 | 1167 | 65.46 |
| Unaltered | 7025 | 2184 | 85.51 |
| ~(PARP and MAP2K) | 10243 | 3351 | 77.65 |

| | coef | exp(coef) | z | p |
|---|---|---|---|---|
| Diagnosis Age | 0.01 | 1.01 | 15.02 | 5.46e-51 |
| Cancer Type | -0.01 | 0.99 | -4.98 | 6.37e-07 |
| Mutation | 0.13 | 1.14 | 3.03 | 2.41e-03 |
| Gender | 0.09 | 1.09 | 4.29 | 1.79e-05 |

Figure S12: **Survival Analysis of simultaneous mutation in PARP and MAP2K groups.** Kaplan Meier survival curve for PARP(1 to 16) and MAP2K(1 to 16) family and log-rank test p-value between the groups: Simultaneous mutation between two groups and the rest of the patients. The median survival time of the group with simultaneous mutation is 18 months more than the rest of the patients. The table at the bottom shows the coxph hazard function where *mutation* parameter is simultaneous mutation between two groups. Although the score of this pair indicates negative pair, we do not have any label for this pair.

##### 3.9.5 DAPK-PARP Survival Analysis on Simultaneous Mutation

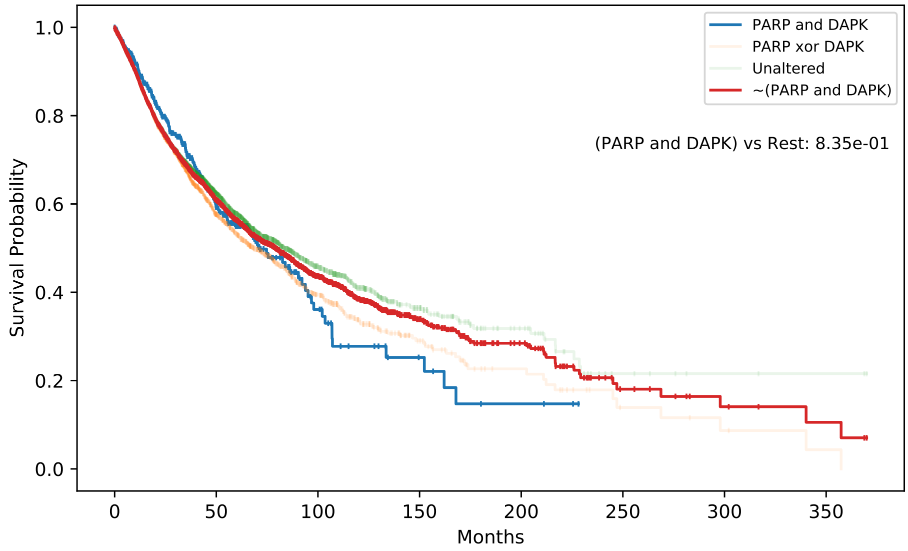

| | # of Cases | # of Events | Median Survival Time |
|---|---|---|---|
| PARP and DAPK | 416 | 147 | 71.01 |
| PARP xor DAPK | 3143 | 1116 | 68.48 |
| Unaltered | 7260 | 2262 | 83.31 |
| ~(PARP and DAPK) | 10403 | 3378 | 79.07 |

| | coef | exp(coef) | z | p |
|---|---|---|---|---|
| Diagnosis Age | 0.01 | 1.01 | 14.94 | 1.92e-50 |
| Cancer Type | -0.01 | 0.99 | -5.05 | 4.38e-07 |
| Mutation | 0.06 | 1.06 | 1.24 | 2.15e-01 |
| Gender | 0.09 | 1.09 | 4.34 | 1.40e-05 |

Figure S13: **Survival Analysis of simultaneous mutation in PARP and DAP2K groups.** Kaplan Meier survival curve for PARP(1 to 16) and DAP2K(1 to 16) family and log-rank test p-value between the groups: Simultaneous mutation between two groups and the rest of the patients. The median survival time of the group with simultaneous mutation is 8 months less than the rest of the patients. The table at the bottom shows the coxph hazard function where *mutation* parameter is simultaneous mutation between two groups. Although the score of this pair indicates negative pair, we do not have any label for this pair.

### 4 Supplementary Tables

Table S1: Number of samples for each cancer type and dataset before removing the duplicates between these datasets, total number of samples for each cancer type after removing disagreeing duplicates and combining datasets. Agree: Duplicates have same label across the datasets they are found. Disagree: Duplicates have different labels across the datasets they are found.

| **Cancer** | **Exp2SL** | | **Lu15** | | **ISLE** | | **dSL** | | **Total** | | **Duplicates** | |
|---|---|---|---|---|---|---|---|---|---|---|---|---|
| | + | - | + | - | + | - | + | - | + | - | **Agree** | **Disagree** |
| BRCA | 0 | 0 | 0 | 0 | 590 | 1012 | 885 | 75 | 1444 | 1037 | 53 | 14 |
| CESC | 0 | 0 | 0 | 0 | 145 | 4762 | 0 | 0 | 145 | 4762 | 0 | 0 |
| COAD | 18 | 155 | 231 | 5621 | 2100 | 74244 | 0 | 0 | 1728 | 79323 | 350 | 484 |
| KIRC | 0 | 0 | 0 | 0 | 60 | 2514 | 0 | 0 | 60 | 2514 | 0 | 0 |
| LAML | 0 | 0 | 0 | 0 | 1191 | 19308 | 0 | 0 | 1191 | 19308 | 0 | 0 |
| LUAD | 307 | 2369 | 0 | 0 | 169 | 4735 | 372 | 339 | 597 | 5515 | 1695 | 242 |
| OV | 0 | 0 | 0 | 0 | 255 | 554 | 0 | 0 | 255 | 554 | 0 | 0 |
| SKCM | 18 | 72 | 0 | 0 | 89 | 18630 | 0 | 0 | 107 | 18702 | 0 | 0 |

Table S2: Number of samples for each cancer in training and testing set after dividing our samples into training(80%) and testing(20%) sets. *Ratio of positive classes* shows $\frac{\#\text{ of positive samples}}{\#\text{ of all samples}}$. It shows that for most of the cancer types, there is a significant imbalance between classes. It also indicates that the random AUPRC performance will be close to these ratios if we do not balance the testing dataset. Since most of them are close to 0, it will be hard to evaluate the AUPRC score.

| **Cancer** | **Train** | | | | **Test** | | | |
|---|---|---|---|---|---|---|---|---|
| | + | - | **Total** | **Pos. ratio** | + | - | **Total** | **Pos. ratio** |
| BRCA | 1155 | 829 | 1984 | 0.582 | 289 | 208 | 497 | 0.581 |
| CESC | 116 | 3809 | 3925 | 0.030 | 29 | 953 | 982 | 0.030 |
| COAD | 1382 | 63458 | 64840 | 0.021 | 346 | 15865 | 16211 | 0.021 |
| KIRC | 48 | 2011 | 2059 | 0.023 | 12 | 503 | 515 | 0.23 |
| LAML | 953 | 15446 | 16399 | 0.058 | 238 | 3862 | 4100 | 0.058 |
| LUAD | 478 | 4411 | 4889 | 0.098 | 119 | 1104 | 1223 | 0.097 |
| OV | 204 | 443 | 647 | 0.315 | 51 | 111 | 162 | 0.315 |
| SKCM | 86 | 14961 | 15047 | 0.006 | 21 | 3741 | 3762 | 0.006 |

Table S3: Train Set Analysis - Top 5 Genes propose a general overview of the genes and their percentage in the positive and negative pairs. The tables show the top 5 genes according to their frequency. *(%)* indicates the percentage of having the gene in pairs. Additionally, $+$ indicates positive pairs, $-$ negative pairs, and the number in the cells indicates the number of times the gene is seen in positive or negative pairs. We can infer from these tables, OV has dense labels while SKCM experiments with only MYC as the first gene in pair. We can also see that the experiments focused on promising genes for each cancer type: PARP1, BRCA1-2, TP53 for BRCA; KRAS for LUAD and COAD; MYC for SKCM. Some genes have only negative or positive relations: BRCA1, PTEN, TP53 is always positively linked in BRCA, while MAP2K1 and HDAC1 are always negatively linked in LUAD.

| **BRCA** | **Count** | (%) | + | - | **CESC** | **Count** | (%) | + | - |
|---|---|---|---|---|---|---|---|---|---|
| PARP1 | 336 | 16 | 42 | 294 | FNTA | 126 | 3 | 4 | 122 |
| BRCA1 | 233 | 11 | 233 | 0 | CHEK2 | 133 | 3 | 4 | 129 |
| PTEN | 202 | 10 | 202 | 0 | HSP90AA1 | 119 | 3 | 3 | 116 |
| TP53 | 131 | 6 | 131 | 0 | HDAC6 | 139 | 3 | 0 | 139 |
| BRCA2 | 88 | 4 | 80 | 8 | HDAC2 | 123 | 3 | 0 | 123 |
| **COAD** | **Count** | (%) | + | - | **KIRC** | **Count** | (%) | + | - |
| KRAS | 14177 | 21 | 113 | 14064 | PIK3CA | 62 | 3 | 0 | 62 |
| PTEN | 11751 | 18 | 325 | 11426 | VHL | 75 | 3 | 2 | 73 |
| BLM | 11597 | 17 | 323 | 11274 | HDAC1 | 63 | 3 | 0 | 63 |
| MUS81 | 11622 | 17 | 326 | 11296 | NF1 | 67 | 3 | 0 | 67 |
| PTTG1 | 11592 | 17 | 275 | 11317 | MET | 60 | 2 | 0 | 60 |
| **LAML** | **Count** | (%) | + | - | **LUAD** | **Count** | (%) | + | - |
| EPHB2 | 166 | 1 | 6 | 160 | KRAS | 571 | 11 | 295 | 276 |
| LARS2 | 174 | 1 | 6 | 168 | CHEK2 | 149 | 3 | 12 | 137 |
| SKP2 | 171 | 1 | 3 | 168 | MAP2K1 | 115 | 2 | 0 | 115 |
| PRKAR1A | 169 | 1 | 8 | 161 | MAPK1 | 103 | 2 | 2 | 101 |
| CYP26A1 | 170 | 1 | 2 | 168 | HDAC1 | 143 | 2 | 0 | 143 |
| **OV** | **Count** | (%) | + | - | **SKCM** | **Count** | (%) | + | - |
| ABL1 | 66 | 10 | 28 | 38 | MYC | 14976 | 99 | 75 | 14901 |
| YES1 | 69 | 10 | 29 | 40 | A1BG | 1 | 0 | 0 | 1 |
| EPHA2 | 66 | 10 | 21 | 45 | POLR3K | 1 | 0 | 0 | 1 |
| KIT | 68 | 10 | 28 | 40 | POM121 | 1 | 0 | 0 | 1 |
| LCK | 67 | 10 | 25 | 42 | POM121C | 1 | 0 | 0 | 1 |

Table S4: Details of the screening experiments that label sources used. It shows the nickname of screening(custom names, not official) if included in specific label sources, the cancer type that cell lines belong to and PMID or doi.

| Screen | ISLE | EXP2SL | LU | dSL | Cancer | PMID or DOI |
|---|---|---|---|---|---|---|
| Zhao | | x | | | CESC, LUAD | 29452643 |
| Big Papi | | x | | | RCC, SKCM, LUAD, COAD, SKCM, OV | 29251726 |
| Han | x | | | | LAML | 28319085 |
| Shen | x | x | | | CESC, LUAD, KIRC | 28319113 |
| ISLE1 | x | | | | | 28162770 |
| ISLE2 | x | | | | CESC | 27453043 |
| ISLE3 | x | | | | | 26612952 |
| ISLE4 | x | | | | OV | 26637171 |
| ISLE5 | x | | | | CESC | 26437225 |
| ISLE6 | x | | | | BRCA | 25407795 |
| ISLE7 | x | | x | | COAD | 24104479 |
| ISLE8 | x | | | | SKCM | 22623531 |
| ISLE9 | x | | | x | KRAS gene | 22613949 |
| ISLE10 | x | | | x | KRAS gene | 19490893 |
| ISLE11 | x | | | | CESC | 20049736 |
| ISLE12 | x | | | x | BRCA | 18388863 |
| ISLE13 | x | | | x | BRCA | 18832051 |
| ISLE14 | x | | | x | KIRC | 18948595 |
| ISLE15 | x | | | | LUAD | 17429401 |
| LU1 | | | x | | COAD | 23563794 |
| dSL1 | | | | x | BRCA, OV, PDAC | 22585861 |
| dSL2 | | | | x | HLRCC | 24568598 |
| dSL3 | | | | x | BRCA, PDAC, OV and UTE | 26427375 |
| dSL4 | | | | x | All | 10.1146/ annurev-cancerbio-042016-073434 |

Table S5: Number of samples used in single cancer and double holdout experiments. Since they are over 10 runs, the first number indicates the mean, and the second shows the standard deviation.

| **Cancer** | **Single Cancer Experiment** | | **Double-Holdout Experiment** | |
|---|---|---|---|---|
| | **Training Samples** | **Testing Samples** | **Training Samples** | **Testing Samples** |
| BRCA | 1658.0 +- 0.0 | 416.0 +- 0.0 | 808.8 +- 113.3 | 239.8 +- 75.4 |
| CESC | 232.0 +- 0.0 | 58.0 +- 0.0 | 116.0 +- 18.4 | 36.0 +- 12.8 |
| COAD | 2764.0 +- 0.0 | 692.0 +- 0.0 | 1571.4 +- 185.1 | 299.8 +- 147.3 |
| KIRC | 96.0 +- 0.0 | 24.0 +- 0.0 | 56.0 +- 12.1 | 12.0 +- 4.8 |
| LAML | 1906.0 +- 0.0 | 476.0 +- 0.0 | 1048.6 +- 75.5 | 255.6 +- 41.4 |
| LUAD | 956.0 +- 0.0 | 238.0 +- 0.0 | 449.8 +- 241.7 | 147.2 +- 95.5 |
| OV | 408.0 +- 0.0 | 102.0 +- 0.0 | 215.0 +- 15.4 | 60.2 +- 9.5 |
| SKCM | 172.0 +- 0.0 | 42.0 +- 0.0 | 61.8 +- 35.8 | 21.0 +- 13.9 |

Table S6: Number of samples used in cross-dataset experiment. Since they are over 10 runs, the first number indicates the mean, and the second shows the standard deviation.

| **Cancer** | **From** | **To** | **Training Samples** | **Testing Samples** |
|---|---|---|---|---|
| BRCA | ISLE | DiscoverSL | 1146.0 +- 0.0 | 150.0 +- 0.0 |
| | DiscoverSL | ISLE | 78.0 +- 0.0 | 1180.0 +- 0.0 |
| LUAD | ISLE | DiscoverSL | 338.0 +- 0.0 | 678.0 +- 0.0 |
| | DiscoverSL | ISLE | 678.0 +- 0.0 | 338.0 +- 0.0 |
| | DiscoverSL | Exp2SL | 678.0 +- 0.0 | 614.0 +- 0.0 |
| | Exp2SL | DiscoverSL | 614.0 +- 0.0 | 678.0 +- 0.0 |

Table S7: The table reports the defaults parameters used in models for the experiments within the training set and fine-tuning set of parameters for the main experiments. These models and parameters are taken from LightGBM[12]

| **Parameter** | **Default value** | **Fine-tuning set** |
|---|---|---|
| Number of leaves | 165 | 165 |
| Max depth | Infinite | [10, 15, 20, ..., 100, 105, 110, Inf] |
| Learning rate | 0.1 | 0.1 |
| No of estimators | 400 | [100, 110, 120, ..., 1180, 1190, 1200] |
| Subsample for bin | 200000 | 200000 |
| Minimum split gain | 0 | 0 |
| Minimum child weight | 5 | 5 |
| Minimum child samples | 10 | {1, 2, 4, 10} |
| Subsample | 0.632 | {0.632, 0.8, 0.99} |
| Subsample frequency | 1 | 1 |
| Colsample by tree | 0.8 | {0.5, 0.8, 1} |
| Alpha regularization | 0 | 0 |
| Lambda regularization | 5 | {5, 10} |

Table S8: Top 10 scored pairs in single cancer experiments for BRCA, LUAD, and OV. For this analysis, we used the ELISL-RF model and mean score over 10 runs. While all top-score predictions have label 1 for BRCA and OV, LUAD has 4 negative and 6 positives in the top 10.

| **Pair Name** | **Score** | **Label** |
|---|---|---|
| | **BRCA** | |
| BRCA1\|THRSP | 0.793051 | 1 |
| BRCA1\|GDNF | 0.789517 | 1 |
| BRCA1\|TSNARE1 | 0.778300 | 1 |
| BRCA1\|SLC7A13 | 0.775389 | 1 |
| BRCA2\|HTR4 | 0.775112 | 1 |
| IL13\|PTEN | 0.771248 | 1 |
| BRCA1\|DNAJC19 | 0.770358 | 1 |
| APOBEC2\|PTEN | 0.768812 | 1 |
| BRCA1\|SLITRK3 | 0.768645 | 1 |
| BRCA1\|ST6GAL1 | 0.763534 | 1 |
| | **LUAD** | |
| KRAS\|MRPL28 | 0.797100 | 0 |
| KRAS\|PARP12 | 0.790102 | 1 |
| KRAS\|LSM5 | 0.789647 | 1 |
| KRAS\|POLR2G | 0.786567 | 1 |
| KRAS\|TEAD2 | 0.783438 | 1 |
| KRAS\|POLL | 0.782135 | 0 |
| KRAS\|MTA2 | 0.779881 | 1 |
| KRAS\|SERPINI1 | 0.778718 | 1 |
| KRAS\|NR1D2 | 0.777780 | 0 |
| KRAS\|OSM | 0.771441 | 0 |
| | **OV** | |
| EPB41L1\|YES1 | 0.640591 | 1 |
| ABL1\|SRC | 0.635870 | 1 |
| FYN\|YES1 | 0.629252 | 1 |
| ABL1\|YES1 | 0.629216 | 1 |
| GAB1\|YES1 | 0.623906 | 1 |
| FYN\|NEDD9 | 0.617254 | 1 |
| ABL1\|BCAR3 | 0.616596 | 1 |
| ABL1\|LCK | 0.614998 | 1 |
| ABL2\|EPB41L1 | 0.604245 | 1 |
| LCK\|PLCG2 | 0.596859 | 1 |

Table S9: Top 50 scored testing and unknown(Unk) pairs in single cancer experiments for BRCA. For this analysis, we used the ELISL-RF model and mean score over 10 runs. The first 38 pairs have a higher score than the top 5% of the positive pair score distribution, 0.757

| Rank | Pair Name | Score | Label | Rank | Pair Name | Score | Label |
|---|---|---|---|---|---|---|---|
| 1 | BRCA1\|THRSP | 0.793051 | 1 | 26 | DBH\|PTEN | 0.761441 | 1 |
| 2 | BRCA1\|GDNF | 0.789517 | 1 | 27 | BRCA2\|NHEJ1 | 0.761429 | Unk |
| 3 | BRCA1\|TSNARE1 | 0.778300 | 1 | 28 | BRCA1\|RPA4 | 0.761338 | Unk |
| 4 | BRCA1\|SLC7A13 | 0.775389 | 1 | 29 | BRCA2\|WNT10A | 0.761283 | Unk |
| 5 | BRCA2\|HTR4 | 0.775112 | 1 | 30 | BRCA1\|FGF19 | 0.760441 | Unk |
| 6 | BRCA1\|HHIP | 0.773132 | Unk | 31 | PTEN\|SFRP5 | 0.760238 | 1 |
| 7 | IL13\|PTEN | 0.771248 | 1 | 32 | BRCA2\|WNT16 | 0.760169 | Unk |
| 8 | BRCA1\|DNAJC19 | 0.770358 | 1 | 33 | BRCA2\|HHIP | 0.759460 | Unk |
| 9 | BRCA2\|FGF6 | 0.769908 | Unk | 34 | BRCA2\|WNT6 | 0.759366 | Unk |
| 10 | BRCA1\|FGF8 | 0.769124 | Unk | 35 | FGF3\|PTEN | 0.758731 | Unk |
| 11 | APOBEC2\|PTEN | 0.768812 | 1 | 36 | BRCA1\|CLDN11 | 0.757921 | 1 |
| 12 | BRCA1\|SLITRK3 | 0.768645 | 1 | 37 | BRCA2\|EPHA5 | 0.757553 | 1 |
| 13 | BRCA1\|WNT3A | 0.767623 | Unk | 38 | BRCA2\|FGF19 | 0.757127 | Unk |
| 14 | BRCA1\|WNT7A | 0.767502 | Unk | 39 | BRCA1\|RAD54L | 0.756919 | Unk |
| 15 | BRCA1\|WNT16 | 0.767323 | Unk | 40 | BRCA1\|EGLN3 | 0.756791 | Unk |
| 16 | BRCA1\|FZD9 | 0.767044 | Unk | 41 | BRCA2\|WNT7B | 0.756601 | Unk |
| 17 | BRCA1\|WNT9A | 0.765874 | Unk | 42 | BRCA1\|FGF11 | 0.755775 | Unk |
| 18 | BRCA1\|FGF6 | 0.765072 | Unk | 43 | OR2T5\|PTEN | 0.755711 | 1 |
| 19 | BRCA1\|NEIL1 | 0.763973 | Unk | 44 | BRCA1\|FASLG | 0.755624 | Unk |
| 20 | BRCA1\|POLD2 | 0.763910 | Unk | 45 | BRCA2\|FGF21 | 0.755194 | Unk |
| 21 | BRCA1\|ST6GAL1 | 0.763534 | 1 | 46 | BRCA1\|WNT7B | 0.755098 | Unk |
| 22 | BRCA1\|FGF21 | 0.763487 | Unk | 47 | BRCA1\|WNT2 | 0.755052 | Unk |
| 23 | BRCA2\|WNT7A | 0.762903 | Unk | 48 | BRCA1\|FATE1 | 0.755044 | 1 |
| 24 | BRCA1\|NEIL2 | 0.762499 | Unk | 49 | BRCA2\|POLD2 | 0.754921 | Unk |
| 25 | BRCA1\|MMP16 | 0.762461 | 1 | 50 | BRCA1\|SRMS | 0.754721 | 1 |

Table S10: The gene sets used and the number of genes for each of them while forming unknown pairs.

| Group | Set Name | Number of Genes |
|---|---|---|
| **CANCER** | KEGG PATHWAYS IN CANCER | 325 |
| **REPAIR PATHWAYS** | KEGG BASE EXCISION REPAIR | 35 |
| | REACTOME BASE EXCISION REPAIR | 91 |
| | WP NUCLEOTIDE EXCISION REPAIR | 44 |
| | KEGG NUCLEOTIDE EXCISION REPAIR | 44 |
| | REACTOME NUCLEOTIDE EXCISION REPAIR | 110 |
| | KEGG MISMATCH REPAIR | 23 |
| | REACTOME MISMATCH REPAIR | 15 |
| | WP DNA MISMATCH REPAIR | 23 |
| | WP HOMOLOGOUS RECOMBINATION | 13 |
| | KEGG HOMOLOGOUS RECOMBINATION | 28 |
| | KEGG NON HOMOLOGOUS END JOINING | 13 |
| | PID FANCONI PATHWAY | 47 |
| **TOTAL** | UNION OF ALL SETS | 572 |

### 5 References

[6] Hussein Hazimeh and Rahul Mazumder. Fast best subset selection: Coordinate descent and local combinatorial optimization algorithms, 2018.

[7] Tim Head, Manoj Kumar, Holger Nahrstaedt, Gilles Louppe, and Iaroslav Shcherbatyi. scikit-optimize/scikit-optimize, September 2020.

Table S11: Survival tables for potential unknown SL pairs: BRCA-HH, BRCA-FGF, BRCA-WNT. Median survival time stands for the time where the half of the population died. For the coxph function, high *coef* (>0.0) and high *exp(coef)* (>1.0) stands for better survival in case of simultaneous mutation. The significance of difference is stated in *p* column. For all three pairs, having simultaneous mutation has an significant positive impact on survival

| **BRCA - HH Survival Table** | | | | **BRCA - HH CoxPH Function** | | | | |
|---|---|---|---|---|---|---|---|---|
| **Group** | **Cases** | **Events** | **Median Survival Time** | **Parameters** | **Coef** | **Exp (Coef)** | **z** | **p** |
| BRCA and HH | 152 | 34 | Inf | **Age** | 0.01 | 1.01 | 14.91 | 2.76e-50 |
| BRCA xor HH | 1256 | 437 | 68.66 | **Cancer** | -0.01 | 0.99 | -4.96 | 7.19e-07 |
| Rest | 9411 | 3054 | 78.44 | **CoMutation** | 0.21 | 1.24 | 2.65 | 8.04e-03 |
| ~(BRCA and HH) | 10667 | 3491 | 77.65 | **Gender** | 0.09 | 1.09 | 4.27 | 1.97e-05 |

| **BRCA - FGF Survival** | | | | **BRCA - FGF CoxPH Function** | | | | |
|---|---|---|---|---|---|---|---|---|
| **Group** | **Cases** | **Events** | **Median Survival Time** | **Parameter** | **Coef** | **Exp (Coef)** | **z** | **p** |
| BRCA and FGF | 451 | 142 | 102.1 | **Age** | 0.01 | 1.01 | 14.98 | 9.89e-51 |
| BRCA xor FGF | 3038 | 1050 | 70.13 | **Cancer** | -0.01 | 0.99 | -5.12 | 3.09e-07 |
| Rest | 7330 | 2333 | 81.2 | **CoMutation** | 0.12 | 1.13 | 2.42 | 1.55e-02 |
| ~(BRCA and FGF) | 10368 | 3383 | 78.44 | **Gender** | 0.09 | 1.09 | 4.35 | 1.39e-05 |

| **BRCA - WNT Survival** | | | | **BRCA - WNT CoxPH Function** | | | | |
|---|---|---|---|---|---|---|---|---|
| **Group** | **Cases** | **Events** | **Median Survival Time** | **Parameters** | **Coef** | **Exp (Coef)** | **z** | **p** |
| BRCA and WNT | 161 | 41 | 167.9 | **Age** | 0.01 | 1.01 | 14.93 | 2.13e-50 |
| BRCA xor WNT | 1291 | 419 | 81.73 | **Cancer** | -0.01 | 0.99 | -5.01 | 5.55e-07 |
| Rest | 9366 | 3065 | 77.19 | **CoMutation** | 0.23 | 1.26 | 2.88 | 3.96e-03 |
| ~(BRCA and WNT) | 10658 | 3483 | 78.21 | **Gender** | 0.09 | 1.09 | 4.32 | 1.56e-05 |

[8] Jiang Huang, Min Wu, Fan Lu, Le Ou-Yang, and Zexuan Zhu. Predicting synthetic lethal interactions in human cancers using graph regularized self-representative matrix factorization. *BMC Bioinformatics*, 20(S19), December 2019.

[9] Jin Huang and C.X. Ling. Using AUC and accuracy in evaluating learning algorithms. *IEEE Transactions on Knowledge and Data Engineering*, 17(3):299–310, March 2005.

[10] J. D. Hunter. Matplotlib: A 2d graphics environment. *Computing in Science & Engineering*, 9(3):90–95, 2007.

[11] L. J. Jensen, M. Kuhn, M. Stark, S. Chaffron, C. Creevey, J. Muller, T. Doerks, P. Julien, A. Roth, M. Simonovic, P. Bork, and C. von Mering. STRING 8–a global view on proteins and their functional interactions in 630 organisms. *Nucleic Acids Research*, 37(Database):D412–D416, January 2009.

[12] Guolin Ke, Qi Meng, Thomas Finley, Taifeng Wang, Wei Chen, Weidong Ma, Qiwei Ye, and Tie-Yan Liu. Lightgbm: A highly efficient gradient boosting decision tree. *Advances in neural information processing systems*, 30:3146–3154, 2017.

[13] Yahui Long, Min Wu, Yong Liu, Jie Zheng, Chee Keong Kwoh, Jiawei Luo, and Xiaoli Li. Graph contextualized attention network for predicting synthetic lethality in human cancers. *Bioinformatics*, February 2021.

[14] B.W. Matthews. Comparison of the predicted and observed secondary structure of t4 phage lysozyme. *Biochimica et Biophysica Acta (BBA) - Protein Structure*, 405(2):442–451, October 1975.

[15] Rose Oughtred, Chris Stark, Bobby-Joe Breitkreutz, Jennifer Rust, Lorrie Boucher, Christie Chang, Nadine Kolas, Lara O'Donnell, Genie Leung, Rochelle McAdam, Frederick Zhang, Sonam Dolma, Andrew Willems, Jasmin Coulombe-Huntington, Andrew Chatr-aryamontri, Kara Dolinski, and Mike Tyers. The BioGRID interaction database: 2019 update. *Nucleic Acids Research*, 47(D1):D529–D541, November 2018.

[16] F. Pedregosa, G. Varoquaux, A. Gramfort, V. Michel, B. Thirion, O. Grisel, M. Blondel, P. Prettenhofer, R. Weiss, V. Dubourg, J. Vanderplas, A. Passos, D. Cournapeau, M. Brucher, M. Perrot, and E. Duchesnay. Scikit-learn: Machine learning in Python. *Journal of Machine Learning Research*, 12:2825–2830, 2011.

[17] Colm Seale, Yasin Tepeli, and Joana P Gonçalves. Overcoming selection bias in synthetic lethality prediction. *Bioinformatics*, 38(18):4360–4368, July 2022.

[18] Lin Shi, Johan A Westerhuis, Johan Rosén, Rikard Landberg, and Carl Brunius. Variable selection and validation in multivariate modelling. *Bioinformatics*, 35(6):972–980, 2019.

[19] Michael L. Waskom. seaborn: statistical data visualization. *Journal of Open Source Software*, 6(60):3021, 2021.
